# Trans-cellular control of synapse properties by a cell type-specific splicing regulator

**DOI:** 10.1101/2022.12.07.519444

**Authors:** Lisa Traunmüller, Jan Schulz, Raul Ortiz, Huijuan Feng, Elisabetta Furlanis, Andrea Gomez, Dietmar Schreiner, Josef Bischofberger, Chaolin Zhang, Peter Scheiffele

## Abstract

The recognition of synaptic partners and specification of synaptic properties are fundamental for the function of neuronal circuits. ‘Terminal selector’ transcription factors coordinate the expression of terminal gene batteries that specify cell type-specific properties. Moreover, pan-neuronal alternative splicing regulators have been implicated in directing neuronal differentiation. However, the cellular logic of how splicing regulators instruct specific synaptic properties remains poorly understood. Here, we combine genome-wide mapping of mRNA targets and cell type-specific loss-of-function studies to uncover the contribution of the nuclear RNA binding protein SLM2 to hippocampal synapse specification. Focusing on hippocampal pyramidal cells and SST-positive GABAergic interneurons, we find that SLM2 preferentially binds and regulates alternative splicing of transcripts encoding synaptic proteins, thereby generating cell type-specific isoforms. In the absence of SLM2, cell type-specification, differentiation, and viability are unaltered and neuronal populations exhibit normal intrinsic properties. By contrast, cell type-specific loss of SLM2 results in highly selective, non-cell autonomous synaptic phenotypes, altered synaptic transmission, and associated defects in a hippocampus-dependent memory task. Thus, alternative splicing provides a critical layer of gene regulation that instructs specification of neuronal connectivity in a trans-synaptic manner.

## Introduction

Neuronal synapses are small but remarkably specialized cell-cell contacts. Across synapses, their strength, the probability of neurotransmitter release, and plasticity properties are tightly controlled and represent the basis for neuronal computations. While individual neuronal cells exhibit reproducible intrinsic properties that are linked to the genetic cell identity, the synaptic properties are a function of both, the pre- and postsynaptic partner cell. Thus, a single neuron can form synapses with dramatically different functional properties on two different target cell types ^1-3^. The genetic mechanisms underlying the specification of these properties are incompletely understood.

Pre- and postsynaptic compartments encompass high concentration of specific protein complexes which coalesce around nascent cell contacts. One candidate mechanism for generating target-specific synapse properties are trans-synaptic recognition codes that recruit select ion channels and neurotransmitter receptors in the opposing synaptic membrane ^4-11^. Post-transcriptional mechanisms such as regulated alternative splicing are hypothesized to play a critical role in this process ^12-14^. Cross-species comparisons demonstrated a significant expansion of alternative exon usage in organisms and tissues with high phenotypic complexity. Thus, alternative splicing programs are particularly complex in the nervous system and have vastly expanded in mammals and primates ^15-17^. Moreover, the high degree of splicing regulation in the brain is accompanied by the expression of a large number of neuronal splicing regulators ^18^. Recent rodent studies mapped developmental and cell type-specific alternative splicing programs in neurons ^14,19-24^. The targets of such regulation are enriched for risk genes associated with neurodevelopmental disorders ^19,25^ and alterations in splicing events are associated with autism spectrum disorders in the human population ^26-28^.

Genetic deletion of pan-neuronal RNA binding proteins results in severe alterations in vast programs of alternative splicing and simultaneous deletion of multiple RBP paralogues often results in embryonic or perinatal lethality ^25,29-33^. These studies firmly established a critical role for alternative splicing regulators in neural development. Loss of such regulators modifies alternative splicing of hundreds of target mRNAs and – in some cases - disrupts cell morphology and viability, accompanied by severe impairments of neuronal function. Thus, it has been difficult to dissociate specific functions of RBPs in controlling synaptic connectivity and function from a more general requirement for cell specification and viability. Other RBPs, such as the KH-domain containing paralogues SLM1 and SLM2 exhibit highly selective expression in neuronal cell types, raising the possibility that they may contribute to the terminal differentiation of these cells ^34-36^. Global genetic ablation of SLM2 results in increased synaptic transmission, loss of long-term potentiation at Schaffer collateral synapses in the hippocampus, and altered animal behavior ^37,38^. However, given the lack of cell type-specific genetic studies the molecular logic of how these neuronal cell type-specific splicing regulators contribute to the acquisition of synaptic properties remains largely unclear. This is further complicated by the fact that a single RBP can control different target exons in different cell populations depending on the cellular context ^31^.

Here, we systematically probed the function of SLM2 which is highly expressed in glutamatergic CA1 and CA3 pyramidal cells and a sub-set of somatostatin-positive GABAergic interneurons in the mouse hippocampus ^35,39^. We tested the hypothesis that SLM2 directly regulates and selectively recruits mRNAs encoding synaptic proteins and, thereby, controls the terminal specification of synaptic properties. We combined genome-wide mapping of SLM2-bound mRNAs *in vivo* with conditional loss-of-function analyses in hippocampal pyramidal cells and SST interneurons in *stratum oriens* of hippocampus area CA1. We find that SLM2 is dispensable for the specification of cell types and their intrinsic properties but selectively controls alternative splicing of synaptic proteins, as well as synaptic function and plasticity in a trans-synaptic manner. We propose that cell type-specific alternative splicing regulators like SLM2 provide a key mechanism for instructing the molecular identity of synaptic interaction modules and neuronal circuit function in mammals.

## Results

### SLM2-bound mRNAs encode synaptic proteins

In the mouse hippocampus, SLM2 is expressed in glutamatergic pyramidal cells but also a sub-population of GABAergic interneurons ^34-36^. These include *oriens-alveus lacunosum-moleculare* (OLM) cells of CA1, a class of somatostatin-positive interneurons. It is unknown whether SLM2 functions are shared between GABAergic and glutamatergic neurons. Within neuronal nuclei SLM2 is concentrated in nuclear sub-structures (Fig.1a). These structures are reminiscent of SAM68 nuclear bodies, membrane-less organelles in non-neuronal cells that require RNAs for their assembly ^40,41^. However, only a fraction of sub-nuclear structures in hippocampal neurons showed SLM2 – SAM68 co-localization (Fig.1b). To identify SLM2-associated RNAs we used eCLIP on mouse whole brain and mouse hippocampal samples. Tag counts obtained with the CTK eCLIP analysis pipeline ^42^ from independent replicates were highly correlated (Table S1, Figure S1a). Consistent with the nuclear localization of SLM2, 77% of the binding events occurred in introns whereas only 2% mapped to exons (Fig.1c). Cross-link induced truncation site (CITS) analysis identified the exact protein-RNA crosslink sites, which are enriched in the UWAA tetramer element (W=U/A; Fig. 1d, Fig,S1b), a motif recognized by SLM2 *in vitro* ^43^. De novo motif discovery using mCross, a computational method to model RBP binding sequence specificity and crosslink sites ^44^, revealed a UUWAAAA 7-mer, with the cross-link occurring at U residues, as the dominant RNA motif bound by endogenous SLM2 *in vivo* (Fig.1e). Furthermore, the UWAA motif is similarly enriched in the whole brain and hippocampal eCLIP data and lacks comparable enrichment when analyzing eCLIP signals from *Slm2*^*KO*^ mice (Fig. S1b-d). High confidence SLM2 binding events in the replicates were identified using CLIPper followed by IDR (Figure 1f, log_2_ fold change ≥2 and FDR? -log_10_≥2, Table S2). Gene ontology analysis of SLM2-bound mRNAs revealed a high enrichment of mRNAs encoding glutamatergic synapse components (Fig.1g, S1e). Amongst the 424 high confidence SLM2 target mRNAs in whole brain samples, 110 were annotated in SynGO ^45^ to encode synaptic proteins, with 59 presynaptic and 49 post-synaptic density components (Fig.1h). These include pan-neuronally expressed mRNAs such as *Nrxn1,2,3, Nlgn1, Lrrtm4, Dlgap1,2, Tenm2* and *Cadm1*, as well as postsynaptic proteins preferentially expressed in GABAergic interneurons such as *Erbb4* and *Gria4* ^46,47^. No significant peaks were observed in size-matched input samples and dense clusters of the UWAA motif in target mRNAs often closely aligned with SLM2 binding events (Fig. 1i). These experiments uncover an array of mRNAs encoding synaptic proteins which are bound by endogenous SLM2 *in vivo*.

**Figure 1.**
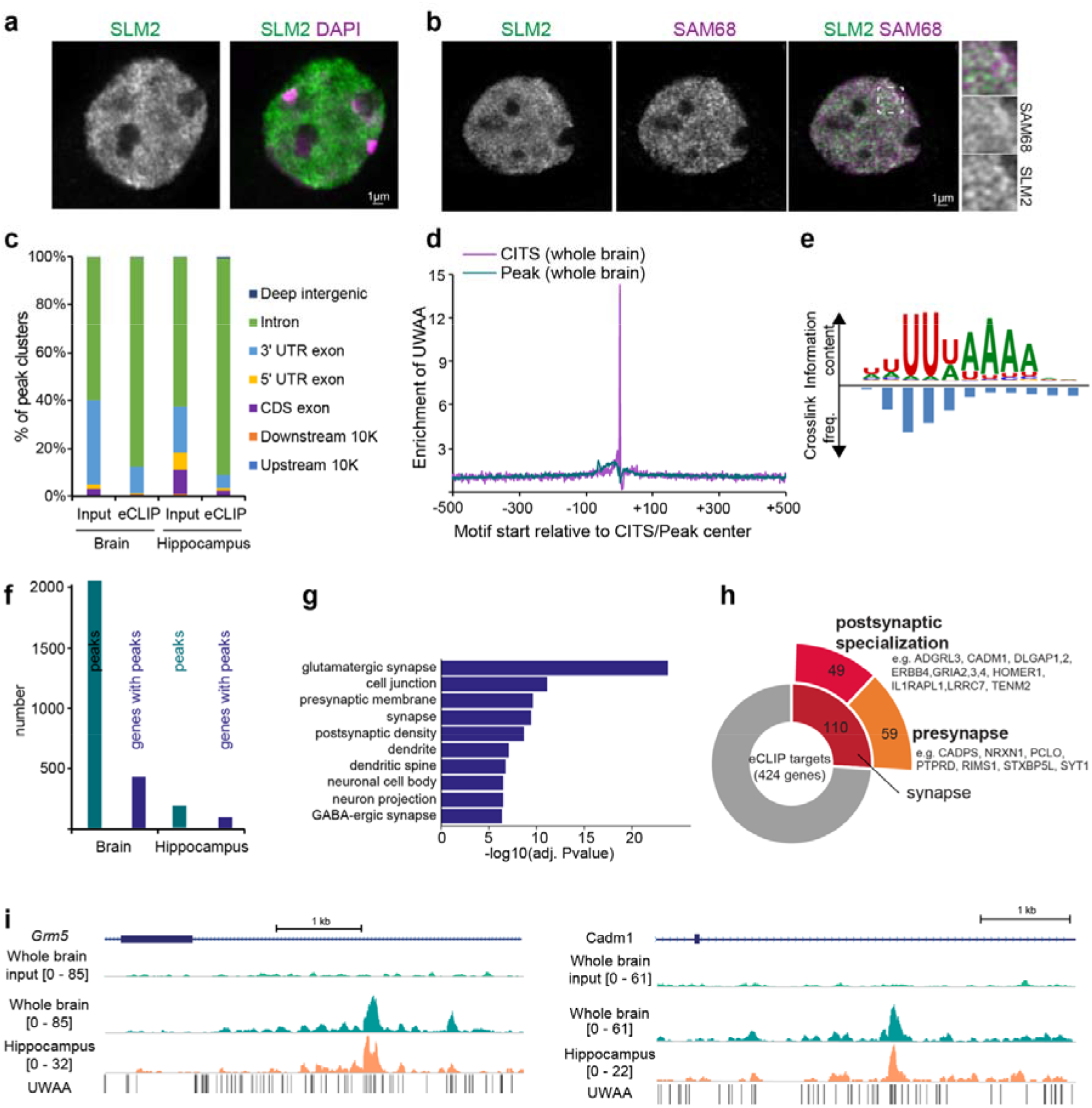
SLM2-bound mRNAs encode synaptic proteins. **a**, Endogenous SLM2 immuno-reactivity in 20µm thick hippocampal sections. DAPI in purple, endogenous SLM2 in green, Scale bar is 1 µm. **b**, Double-immunostaining for SLM2 (green) and SAM68 (magenta) in cultured hippocampal neurons maintained in vitro for 12 days. Insets on the right show an enlargement of the area boxed in the merged image. Scale bar 1µm. **c**, eCLIP cluster annotation summary for input and SLM2 eCLIP samples from whole brain and hippocampus. **d**, Enrichment of UWAA around cross-linked induced truncation site (CITS), calculated from the frequency of UWAA starting at each position relative to the inferred crosslink sites, normalized by the frequency of the element in flanking sequences in whole brain eCLIP data. Enrichment of UWAA around the CLIP tag cluster peak center is shown for comparison. **e**, SLM2 binding motif determined by mCross based on whole brain eCLIP data. Corresponding crosslinking frequencies at each motif position are represented by the blue bars. **f**, Number of high confidence SLM2 targets identified by CLIPper and IDR analysis (log_2_ fold change ≥2 and -log_10_(IDR)≥2) in whole brain and hippocampal eCLIP samples. Note that the lower number of peaks and genes with peaks for hippocampal samples is likely due to the lower read counts obtained for eCLIP libraries generated from limited starting material. **g**, Gene Ontology analysis (DAVID tools) of genes with significant SLM2 binding sites (whole brain). Top 10 enriched gene ontology categories for cellular compartment are displayed. **h**, Sunburst chart and gene examples associated with synaptic function of high confidence eCLIP targets from whole brain samples identified by CLIPper and IDR. **i** SLM2 eCLIP read densities on immunoprecipitated samples show strong enrichment on identified eCLIP targets as compared to size-matched input samples. UWAA motif enrichment of whole brain (green) and hippocampal (orange) samples in introns of the glutamate metabotropic receptor 5 (*Grm5*) and the synaptic cell adhesion molecule *Cadm1* (SynCAM1). Coordinates shown are *Grm5* chr7: 87,601,936-87,607,378 and *Cadm1* chr9: 47,836,291-47,840,954.

### Identification of cell type-specific SLM2-dependent exons

Considering that action of RBPs is frequently graded, i.e. dependent on expression level ^22,48^, we quantified SLM2 immune-reactivity across genetically marked hippocampal neuron subpopulations (Fig.2a,b). We found that SLM2 expression was highest in CA3 pyramidal cells (marked by Grik4-cre, see Fig. S2a) and significantly expressed in CA1 pyramidal cells (CamK2-cre) and SST interneurons (SST-cre) of area CA1 and CA3 (Fig.2b and Fig. S2b). More than 90% of genetically marked CA1 and CA3 pyramidal cells expressed SLM2 (Fig. S2c,d). By contrast, SLM2 is not detectable in dentate granule cells (Fig. 2b). To define SLM2-dependent mRNA splicing events in GABAergic versus glutamatergic populations, we performed conditional ablation and mapped ribosome-associated transcripts in the respective cell types by RiboTrap ^49,50^. Using CamK2-cre, Grik4-cre, and SST-cre lines we selectively ablated SLM2 in hippocampal CA1 (*Slm2*^*ΔCamK2*^) and CA3 (*Slm2*^*ΔGrik4*^) pyramidal cells, and somatostatin-positive GABAergic interneurons (*Slm2*^*ΔSST*^), respectively (Fig.2c,d). All resulting conditional knock-out mice were viable and fertile and did not show overt physical alterations. Immunostaining for SLM2 confirmed complete loss of the protein at postnatal day 16-18 (p16-18) for *Slm2*^*ΔSST*^ and p42-45 for *Slm2*^*ΔCamK2*^ and *Slm2*^*ΔGrik4*^ in 75-90% of the cre-positive cells (Fig.2d). Using cell type-specific RiboTrap affinity isolation of actively translated mRNAs we deeply mapped the transcriptomes in wild-type and conditional knock-out cells (>90 Mio uniquely mapping reads/sample, >84% of reads mapping to mRNA, 4 replicates per genotype and cell population, one replicate for *Slm2*^*ΔCamK2*^ excluded due to 3’ bias, see Table S3 for details). Principal component analysis of the resulting datasets uncovered highly similar transcriptomes within the respective cell type-preparations (Fig.2e). There was very little variance between replicates or the RiboTrap samples from knock-out versus wild-type mice, suggesting that loss of SLM2 does not impact the terminal gene batteries of these cell types. Loss of RNA-binding proteins can exhibit broad effects on transcript abundance and stability. However, scatterplots further confirmed only minimal alterations at the level of overall gene expression between *Slm2* conditional knock-out and wild-type cells (Fig.2f) with only 1-5 genes exhibiting statistically significant differences in expression (adj. p-value ≤0.05 and fold change ≥ 1.5). The most strongly altered transcript (1.5-4.7fold up-regulated, see Table S3 for details) was the SLM2 paralogue SLM1/*Khdrbs2*, consistent with the functional cross-repression between SLM1/2 paralogues ^36^.

**Figure 2.**
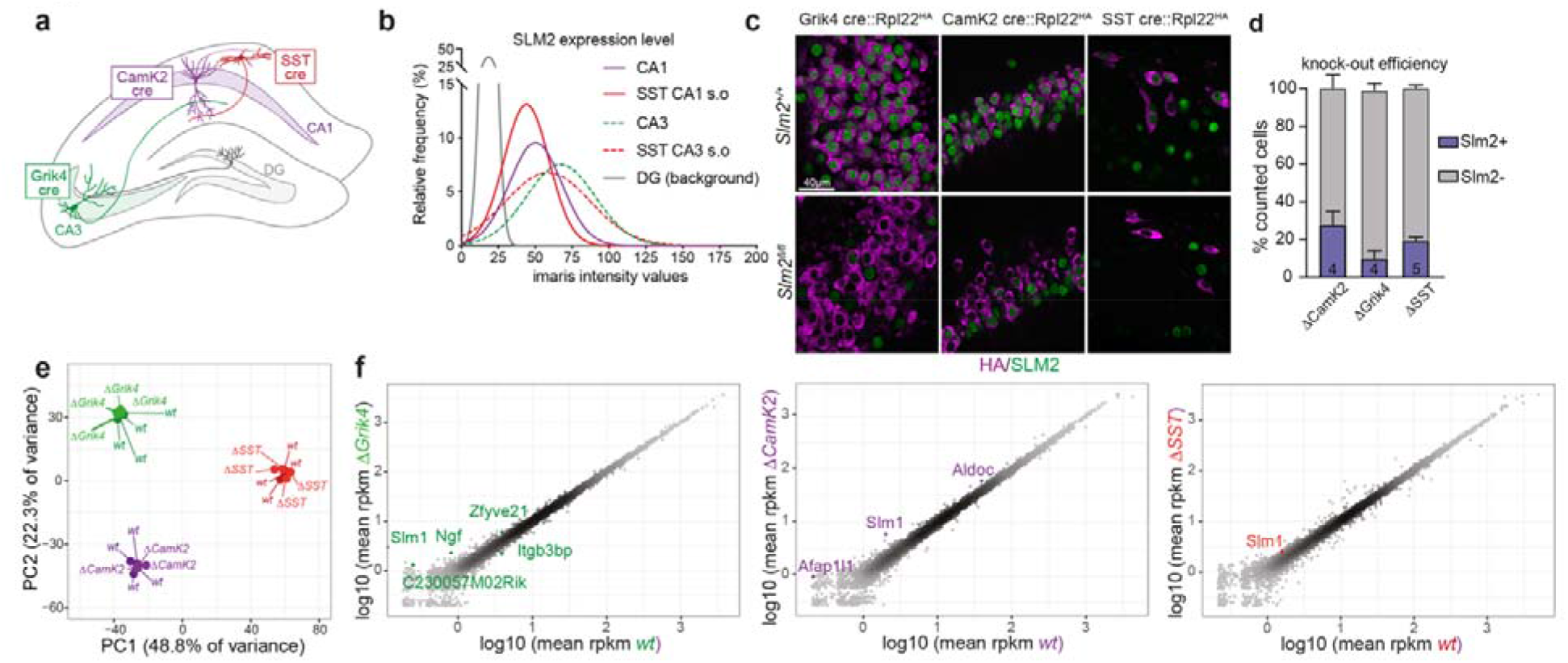
Conditional ablation of SLM2 in hippocampal cell types. **a**, Schematic representation of cre drivers used to assess the molecular profile of hippocampal *Cornu Ammonis* (CA) 1 (CamK2-cre), CA3 (Grik4-cre) and SST-positive (SST-cre) neurons. **b**, Fitted gaussian curves of relative frequency of SLM2 immune-reactivity in genetically marked CA1, CA3 and SST cre-positive neurons in the *stratum oriens* (s.o) of CA1 and CA3. The level of “background” immune-reactivity observed in dentate granule (DG) cells was defined based on comparisons with antibody staining of global *Slm2*^*KO*^ mice. N=3 animals each. CA1: 68 cells, CA3: 75 cells, SST CA1 s.o: 73 cells, SST CA3 s.o: 60 cells, DG: 62 cells. **c**, Representative images of SLM2 expression in cre-positive cells defined by immune-reactivity for the conditional HA-tagged Rpl22 allele in *Slm2*^*+/+*^ and *Slm2*^*fl/fl*^ mice (HA in magenta, SLM2 in green, scale bar 40µm). **d**, Quantification of SLM2 knock-out efficiency in postnatal day (p) 42-45 CA1 (ΔCamK2, N=4, n=1081) and CA3 (ΔGrik4, N=4, n=1070), and p16-18 SST (ΔSST, N=5, n=157) neurons. Displayed as mean ± SEM. SLM2-levels were based on background levels of antibody staining in global *Slm2*^*KO*^ mice. **e**, Principal component analysis of genes expressed in hippocampal *Slm2*-wild-type and conditional knockout RiboTRAP pulldowns (*wt* in green: Grik4^cre^::Rpl22^HA/HA^ N=4, Δ*Grik4*: *Grik4*^*cre*^*::Rpl22*^*HA/HA*^*::Slm2*^*fl/fl*^ N=4, *wt* in purple: *Camk2*^*cre*^*::Rpl22*^*HA/HA*^ N=3, Δ*CamK2*: *Camk2*^*cre*^*::Rpl22*^*HA/HA*^*::Slm2*^*fl/fl*^ N=3, *wt* in red: *SST*^*cre*^*::Rpl22*^*HA/HA*^ N=4, Δ*SST*: *SST*^*cre*^*::Rpl22*^*HA/HA*^*::Slm2*^*fl/fl*^ N=4). Variances explained by principal component 1 (PC1) and 2 (PC2) are indicated. Variance stabilization transformation was utilized to normalize gene expression. **f**, Correlation analysis of the mean log10 transformed, normalized transcript counts (reads per kilobase million, rpkm) between *wt* (x-Axis) and *Slm2*^*D*^ conditional mutants (y-Axis). Significant differentially expressed genes (fold change ≥1.5, adjusted p-value Benjamini and Hochberg ≤0.05) are marked in green for comparisons in RiboTrap pulldowns for CA3, purple for CA1 and red for SST.

We then performed genome-wide mapping of alternative splicing changes elicited by the conditional loss of SLM2. When comparing differential alternative exon usage across wild-type CamK2, Grik4, and SST cells, we identified 2860 differentially regulated exons between these populations (Table S4). Loss of SLM2 did not broadly modify these cell type-specific splicing signatures (Fig.S3a and Table S4). Instead, conditional SLM2 knock-out resulted in significant de-regulation of only a handful of alternative splicing events (Fig.3a, p-value ≤0.01, fold change ≥ 2, Table S4, S5). Notably, loss of SLM2 resulted in increased exon incorporation at *Nrxn2* alternatively spliced segment 4 in all three cell populations (AS4, Fig.3a,b and Fig. S3e for validation of splicing changes by qPCR, and Fig. S4a for a sashimi plot). Thus, all cell populations exhibited a high degree of exon skipping at *Nrxn2* AS4 in wild-type mice but exon mis-incorporation in the conditional knock-outs. By contrast, the corresponding alternative exon in *Nrxn3* was de-regulated only in CA3 (Grik4) and CA1 (CamK2) cells but was not SLM2-dependent in SST interneurons (Fig.3c, Fig.S3e). This reveals significant cell type-specific differences in SLM2-dependent alternative splicing regulation. De-regulation of the mutually exclusive alternative exons e23/e24 in Syntaxin binding protein 5-like (*Stxbp5l*, also called Tomosyn-2) was another splicing event commonly altered in CA1 and CA3 but not SST *Slm2* conditional knock-out neurons (Fig.3a, S3c). In addition, our computational pipeline identified de-regulation of alternative exons in the unconventional myosin 1b (*Myo1b*) (in *Slm2*^*ΔCamk2*^ cells) and the ubiquitin ligase *Ube2d* and the GTPase-activating enzyme *Arhgap42* in *Slm2*^*ΔSST*^ cells.

Integration of the eCLIP and RiboTrap splicing analysis uncovered densely clustered intronic SLM2 binding events and repeats of the UWAA motif within 500 bases downstream of the de-regulated alternative exons of *Nrxn1,2,3* and *Stxbp5l* (Fig.3c). This demonstrates that SLM2 binding directs skipping of upstream alternative exons in the endogenous mRNA. Importantly, no significant eCLIP tags were recovered in *Myo1b, Nkd2, Ube2d*, and *Arhgap42* indicating that alternative splicing of these mRNAs is not directly regulated by SLM2. Besides these major alterations in a handful of genes, we observed further alterations in alternative exon incorporation in 61 additional mRNAs. Notably, the vast majority of these mRNAs are only very lowly expressed (Fig.S3b), indicating that the mRNAs are unlikely to have significant contribution to the cellular proteomes. Moreover, no eCLIP binding events were mapped to these mRNAs demonstrating that they are not directly bound by endogenous SLM2 (Fig. S4b for a Venn diagram summarizing this data). Notably, all directly bound mRNAs with significantly altered alternative splicing in the conditional knock-out cells, encode synaptic proteins. This strongly supports the hypothesis that SLM2 specifically controls synaptic properties in the mouse hippocampus in vivo.

### Loss of SLM2 results in cell type-specific synaptic phenotypes

Considering the remarkable selectivity of SLM2 for binding and regulating mRNAs encoding synaptic proteins we probed the functional consequences of conditional SLM2 ablation in the hippocampal circuit. Global ablation of SLM2 is accompanied by increased evoked glutamatergic transmission at CA3-CA1 pyramidal cell Schaffer collateral synapses and increased postsynaptic AMPA-receptor accumulation ^37^. However, it is unknown whether this phenotype arises from loss of SLM2 in the presynaptic CA3 cells and disruption of trans-synaptic interactions or whether it involves other cell types. Notably, conditional ablation of SLM2 in the presynaptic CA3 pyramidal cells resulted in a significant increase in postsynaptic currents evoked by Schaffer collateral stimulation in CA1 neurons (Fig.S5a-c). Thus, deletion of SLM2 from CA3 neurons is sufficient to modify synaptic transmission onto postsynaptic CA1 pyramidal cells.

We next examined phenotypes resulting from conditional loss of SLM2 in GABAergic interneurons. We focused on horizontally oriented somatostatin interneurons located in the *stratum oriens alveus* of the hippocampus representing putative OLM interneurons which significantly express SLM2 (Fig.S2b). These dendrite-targeting inhibitory interneurons shape hippocampal information processing by gating excitatory transmission and associative synaptic plasticity in CA1 pyramidal cells ^51,52^. Conditional knock-out of SLM2 from SST interneurons (*Slm2*^*ΔSST*^) did not modify resting membrane potential, excitability or other intrinsic properties of SST+ interneurons, indicating that SLM2 is not required for the specification of these cells (Fig.S5c-f). Our eCLIP analysis uncovered abundant SLM2 binding to mRNAs that encode proteins of glutamatergic synapses (Fig.1g,h). Thus, we examined glutamatergic inputs to *Slm2*^*ΔSST*^ cells. mEPSC amplitudes in SST interneurons were unchanged but we observed a significant shift towards a higher mEPSC frequency, suggesting an increased glutamatergic synapse density onto *Slm2*^*ΔSST*^ cells (Fig.4a-e). OLM interneuron dendrites in the *stratum oriens* receive glutamatergic synapses from CA1 pyramidal cells ^51^. These inputs exhibit a characteristic short-term facilitation which has a critical impact on hippocampal circuit function ^53,54^. We investigated AMPAR-mediated post-synaptic responses with increasing electrical stimulation of putative CA1 axons in the *alveus* and found a significant increase in excitation consistent with a larger synapse number (Fig. 4f). Moreover, 40 Hz stimulation of the same axons led to a significantly elevated short-term facilitation (Fig 4g). Because short-term facilitation at this synapse is mediated via increased transmitter release, these results show that SLM2 in postsynaptic SST interneurons controls glutamatergic transmission and synaptic recruitment of these cells via a trans-synaptic mechanism.

**Figure 3.**
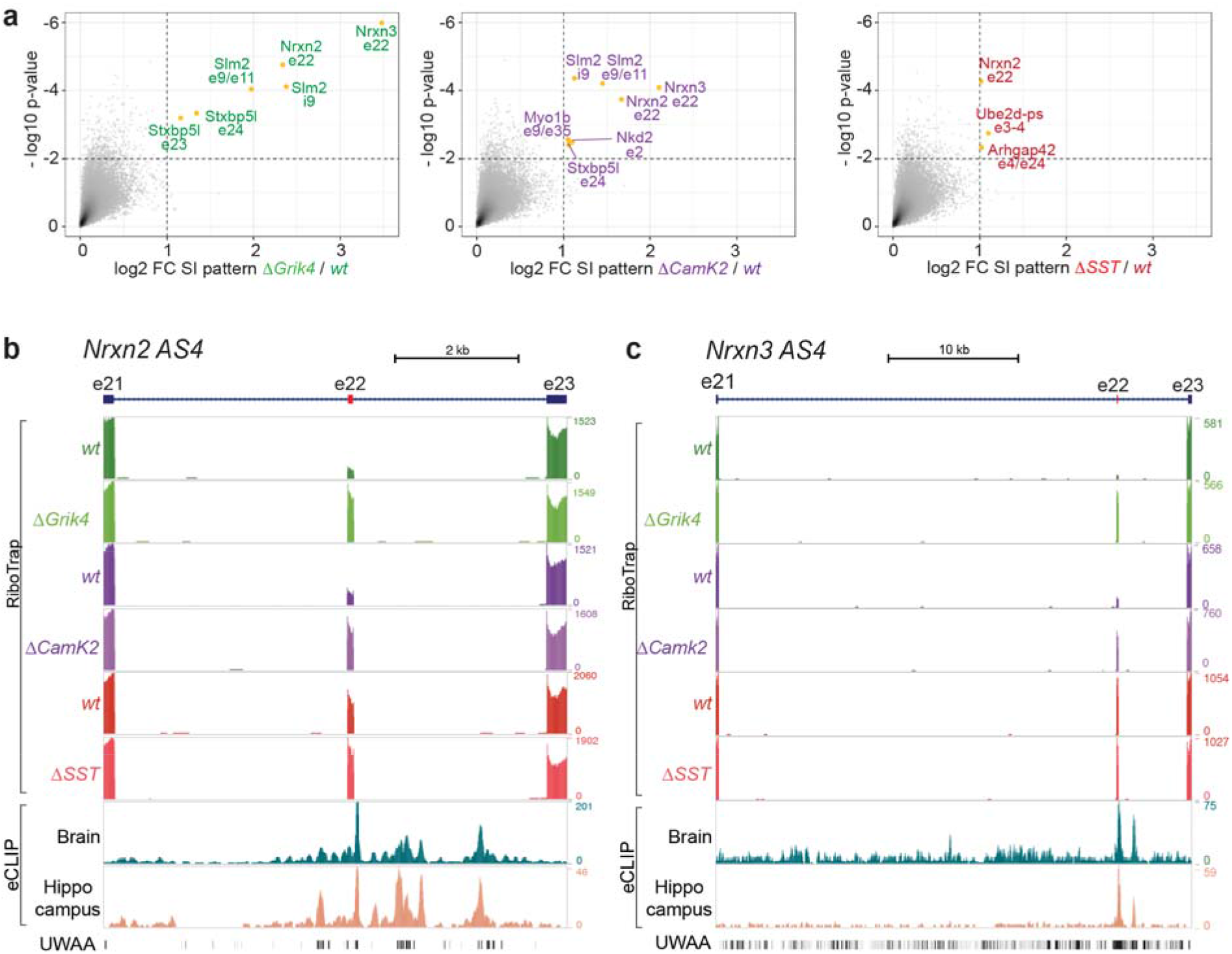
SLM2 directly regulates alternative splicing of mRNAs encoding synaptic proteins. **a**, Log2 fold change splicing index (FC SI) and -Log10 p-values of all detected splicing patterns comparing *wt* and *Slm2* conditional mutants in the different RiboTrap preparations. Significantly differentially regulated exons are marked in green for Grik4, purple for CamK2 and red for SST (Fold change ≥ 2, p-value ≤ 0.01). **b**,**c**, Integration of sequencing tracks for wildtype (wt) and mutant (Δ) RiboTrap samples for hippocampal Grik4, CamK2, and SST cells and eCLIP analysis in whole brain and hippocampus for significantly de-regulated exons of *Nrxn2* and *Nrxn3*. Clusters of SLM2 binding events in downstream introns align with UWAA binding motifs. Coordinates shown are: *Nrxn2* chr19:6,509,778-6,517,248; *Nrxn3* chr12:90,168,920-90,204,818

**Figure 4.**
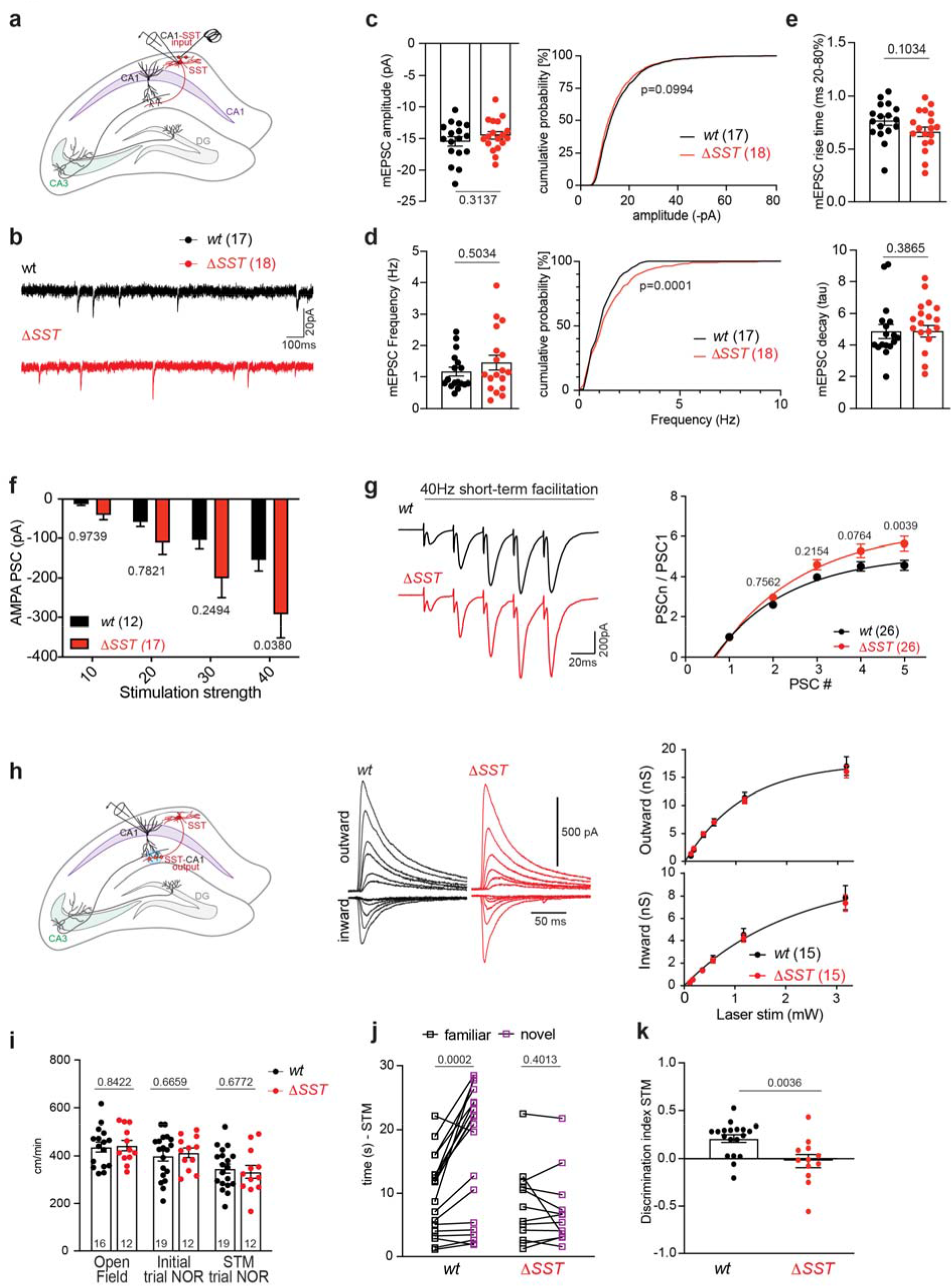
SLM2 controls synaptic plasticity and function in a trans-synaptic manner. **a**, Experimental design for electrical stimulation of axons in the alveus and voltage clamp recordings in genetically marked SST+ interneurons in the *stratum oriens*. **b**, Example traces of miniature excitatory post-synaptic currents (mEPSCs) in SST interneurons. **c**,**d**,**e**, Analysis of mEPSC amplitude (c), frequency (d), rise and decay times (e). *wt* n= 17, Δ*SST* n=18. Mean ± SEM. P-values were determined by the corresponding t-tests based on assessment of normality distribution and standard deviation. For cumulative frequency distribution the Kolmogorov-Smirnov test was performed. **f**, Mean ± SEM data of post-synaptic AMPAR-mediated EPSCs in SST-interneurons of the stratum oriens in response to different stimulation intensities of axons in the alveus is increased in Δ*SST* n=17 versus *wt* n= 12. Two-way ANOVA with Šídák’s multiple comparisons test. **g**, Representative traces of evoked EPSCs during repetitive stimulation at 40Hz, *wt* (black) and Δ*SST* (red). Group data of EPSCs normalized to the first peak show increased facilitation in Δ*SST* n=26 versus *wt* n=26. Mean ± SEM. Two-way ANOVA with Šídák’s multiple comparisons test. **h**, Experimental design for local optogenetic stimulation of SST+ neuron-mediated IPSCs in apical dendrites of CA1 pyramidal neurons in *wt* and Δ*SST* mutants. Representative traces of inward and outward IPSCs evoked at increasing laser intensities in *wt* (black) and Δ*SST* (red) mutants. Mean ±SEM input-output curve of synaptic conductance underlying outward and inward PSCs. **i**, Quantification of velocity (cm/min) of mice during the open field, initial, and short-term memory (STM) test phases of the novel object recognition task. Animal numbers for each task are indicated, Mean ± SEM, Unpaired t-test. **j**,**k**, Behavioral alterations in wt and Δ*SST* animals in the novel object recognition task. Interaction time (in seconds) that mice spend with either a familiar (black) or novel (purple) object during a 5-min short-term memory trial (paired t-test) and discrimination index (unpaired t-test) are displayed. Mean ± SEM, *wt* n=19 and Δ*SST* n=12.

We further analyzed GABAergic SST interneuron output synapses onto CA1. We used optogenetic stimulation of SST interneurons and performed whole-cell patch-clamp recordings from CA1 pyramidal cells in acute hippocampal brain slices. We found no alterations in the magnitude of optogenetically-evoked postsynaptic inhibitory currents in CA1 neurons from *Slm2*^*ΔSST*^ mice (Fig.4h). Moreover, the kinetics of optogenetically evoked currents were unchanged, indicating normal assembly of postsynaptic GABA A receptors in CA1 neurons (Fig.S5g). To assess short-term plasticity of evoked transmission, we applied 10 Hz optogenetic stimulation which induces a depression at OLM-CA1 synapses in wild-type cells. Using this protocol, we observed a small but significant reduction in short-term depression in slices from *Slm2*^*ΔSST*^ mice (Fig.S5h). Finally, GABA A receptor kinetics and voltage-dependence of GABAergic IPSCs were unchanged (Fig. S5), suggesting that expression of synaptic GABA A receptor subunits is virtually identical. Thus, selective loss of SLM2 from SST interneurons results in increased glutamatergic drive onto OLM-interneurons with largely similar properties of output synapses onto CA1 neurons.

Aberrant activation of OLM interneurons induced by optogenetic stimulation during the exploration phase has been shown to impair object memory in an object recognition task ^55^. To test whether the increased glutamatergic drive onto SLM2-deficient SST interneurons is associated with memory deficits we performed novel object recognition tests with *Slm2*^*ΔSST*^ mice. Mutant and wild-type mice did not differ in mobility in the test arena or the total time spent interacting with objects (Fig.4i). When testing object recognition memory (1 hr after the initial object exploration), wild-type mice spent significantly more time exploring the novel object. By contrast, *Slm2*^*ΔSST*^ mice spent similar times interacting with novel and familiar objects (Fig.4j,k). This defect in short term memory was not associated with an increase in anxiety as *Slm2*^*ΔSST*^ mice showed normal exploration of open and closed arms in elevated plus maze and also did not differ in other behavioral assessments such as marble burying (Fig.S6). Thus, selective loss of SLM2 from SST-interneurons is associated with a specific deficit in short-term memory in mice. Taken together, these data suggest, that in SST interneurons, SLM2 controls splicing of a very small subset of mRNAs encoding synaptic proteins including *Nrxn2*, leading to regulation of the glutamatergic recruitment of these GABAergic interneurons for fine-tuning of dendritic inhibition during learning and memory.

## Discussion

Alternative splicing has emerged as a critical and widespread gene regulatory mechanism across organisms and tissues. In the nervous system, alternative splicing controls multiple steps of neuronal development, plasticity, and diverse pathologies ^12,14,16,56^. Individual splicing regulators can govern neuronal viability ^57^, cell fate ^19^, axon guidance ^58^, and broader aspects of neuronal function. In this work, we discovered that SLM2, an RNA binding protein exhibiting highly selective cell type-specific expression, is dispensable for most aspects of neuronal differentiation but selectively instructs terminal specification of synaptic function within the hippocampal microcircuit.

*In vitro* studies identified SLM2 RNA binding motifs as U(U/A)AA repeats ^43,59^. Using eCLIP, we now demonstrate that endogenous SLM2 binds to a UUWAAAA 7-mer motif in vivo. The SLM2 paralogue SAM68 recognizes a similar motif in vitro. However, a direct comparison of alternative splicing profiles in *Slm2*^*KO*^ and *Sam68*^*KO*^ hippocampi suggests that de-regulated exons are largely paralogue-specific *in vivo* (Figure S4c). The endogenous SLM2 eCLIP targets are strongly enriched for mRNAs encoding synaptic proteins, including adhesion molecules, pre- and postsynaptic scaffolding molecules, and neurotransmitter receptors. Interestingly, only a small fraction of these SLM2-bound mRNAs exhibits alterations in alternative exon incorporation in conditional knockout mice. This might be a consequence of functional redundancy with other RBPs. Alternatively, SLM2-binding to target mRNAs in the nucleus may contribute to coordinated spatio-temporal control of an array of functionally related mRNAs which modifies their trafficking and/or translation ^60^. Notably, for all regulated alternative exons, SLM2 binding sites consist of extended RNA motif clusters positioned in the downstream intron. Thus, clustered motifs are a pre-requisite for splicing regulation by SLM2 *in vivo*. In all cases, loss of SLM2 results in aberrant exon incorporation, indicating a major role for SLM2 in driving exon skipping in wild-type cells.

Interestingly, loss of SLM2 did not result in significant alterations in the overall neuronal transcriptomes or functional intrinsic properties of SST-interneurons. Similarly, overall mRNA expression was unaltered in CA1 and CA3 pyramidal cells. This strongly suggests that SLM2 is dispensable for cell fate specification and the acquisition of functional properties of these cells. By contrast, SLM2 loss-of-function was associated with selective trans-synaptic phenotypes: Conditional ablation of SLM2 from CA3 pyramidal neurons led to an increase in postsynaptic currents at Schaffer collateral synapses onto CA1. This phenotype recapitulates the increase in postsynaptic AMPA receptors and increased synaptic transmission observed in global *Slm2*^*KO*^ mice ^37^. In SST-interneurons, conditional *Slm2* deletion was associated with increased glutamatergic transmission likely resulting from an increased glutamatergic synapse density onto the mutant cells and increased presynaptic facilitation of synapses formed onto the knock-out cells. SLM2-dependent alterations in synaptic adhesion molecules in SST interneurons, such as TENM2, ERBB4, CADM1, ADGRL3, and postsynaptic NRXNs are well-positioned to direct such trans-synaptic regulation ^46,61,62^. For example, the elevated alternative exon incorporation in NRXN2 AS4 is predicted to reduce its ability to inactivate the function of postsynaptic neuroligins in neuronal dendrites ^63,64^ and may contribute to the increased glutamatergic input received by OLM interneurons in *Slm2*^*ΔSST*^ mice. Thus, conditional SLM2 ablation from either CA3 or SST neurons does not significantly modify intrinsic properties of the cell populations themselves but reconfigures trans-synaptic interaction modules and, thereby, properties of synaptic structures formed with connecting cells.

Despite the increased glutamatergic drive received by OLM interneurons, their GABAergic output was largely unchanged in *Slm2*^*ΔSST*^ mice. This suggests that SLM2 regulates the level of functional recruitment of SST interneurons via altered splicing of a small set of mRNAs encoding synaptic proteins. Thereby, the RNA binding protein SLM2 shapes activation of CA1 cell assemblies and hippocampal information processing. SST interneurons provide branch-specific inhibition onto distal dendrites of CA1 pyramidal cells, powerfully controlling dendritic integration of synaptic information ^65^. Increased activation of OLM interneurons during the formation of episodic memories has been shown to disrupt memory formation ^55^. Moreover, short-term facilitation (which is altered in *Slm2*^*ΔSST*^ mice) is thought to be involved in the short-term storage of information ^66^. Consistent with an aberrant activation of OLM-interneurons, SLM2 knockout mice exhibit an impairment in short-term memory in the hippocampus-dependent novel object recognition task. At the same time, other behaviors were indistinguishable from wildtype mice. While we cannot exclude a contribution of SST-interneuron populations besides OLM cells to the behavioral phenotype, our results support a critical function for SLM2 in the inhibitory control of short-term episodic memories.

We propose that acquisition of a cell type-specific complement of RNA binding proteins represents a critical element of the terminal gene battery established during development which shapes trans-synaptic modules ^4^. Expression of SLM2 in a sub-class of SST interneurons is already detected at embryonic stages while cells are migrating towards their final cortical locations ^67^. Similarly, we observed expression of SLM2 in hippocampal pyramidal cells already at embryonic stages (Figure S7). Thus, expression of this regulator is indeed temporally linked to embryonic cell type specification. Evolutionary comparisons of synaptic building blocks across organisms suggest that more complex cellular modules accommodate the need for phenotypic diversity at the level of individual synapses ^68^. Our work suggests that the modification of synaptic modules through alternative splicing is a major mechanism underlying the unique functional specification of synaptic connections.

## Limitations of the study

While our study correlates alterations in alternative splicing and synaptic transmission phenotypes we have not directly linked a single alternative splice isoform of a synaptic protein to the alteration in plasticity. While we explored electrophysiological phenotypes resulting from conditional SLM2-ablation in CA3 and SST neurons, we did not examine the consequences of SLM2 deletion by the CamK2-cre driver in CA1 neurons, as the knock-out was only partial and, thus, results would be difficult to interpret.

## Acknowledgements

We thank Caroline Bornmann and Sabrina Innocenti for organizational and experimental support, Geoffrey Fucile (SciCORE) for help with data analysis, the Biozentrum Imaging Core Facility for support with image acquisition and analysis, and the Quantitative Genomics Centre of the University of Basel for excellent technical assistance. This work was supported by funds to P.S. from the Canton Basel-Stadt/University of Basel, the Swiss National Science Foundation (project 179432), a Swiss National Science Foundation Advanced Grant (TMAG-3-209273), a European Research Council Advanced Grant (SPLICECODE), and AIMS-2-TRIALS which are supported by the Innovative Medicines Initiatives from the European Commission, funds to J.B (SNSF, Project 176321) and C.Z. (National Institutes of Health, R01NS125018).

## Supplementary Material

### Supplementary Figures

Figure S1 – related to Figure 1

Figure S2 – related to Figure 2

Figure S3 – related to Figure 3

Figure S4 – related to Figure 4

Figure S5 – related to Figure 4

Figure S6 - related to Figure 4

Figure S7 - related to Figure 2

### Supplementary Tables

**Table S1**. eCLIP data by CTK pipeline for SLM2-bound mRNAs in mouse brain and hippocampal tissue. Tabs list annotation of all clusters identified in whole brain and hippocampus, respectively, called using the CTK pipeline ^44^.

**Tabl2 S2**. eCLIP data by CLIPper/IDR pipeline for SLM2-bound mRNAs in mouse brain and hippocampal tissue.

**Table S3**. Global transcriptome analysis of hippocampal wild type and Slm2 conditional knockout.

**Table S4**. Cell type specific alternative exons differentially regulated between wild type CA1, CA3 and SST interneurons in the hippocampus.

**Table S5**. Alternative exons differentially expressed between *Slm2* control and *Slm2* conditional knockout.

**Table S6**. Differentially regulated patterns of alternative splicing in *Slm2* control and *Slm2* conditional knockout.

**Figure S1.**
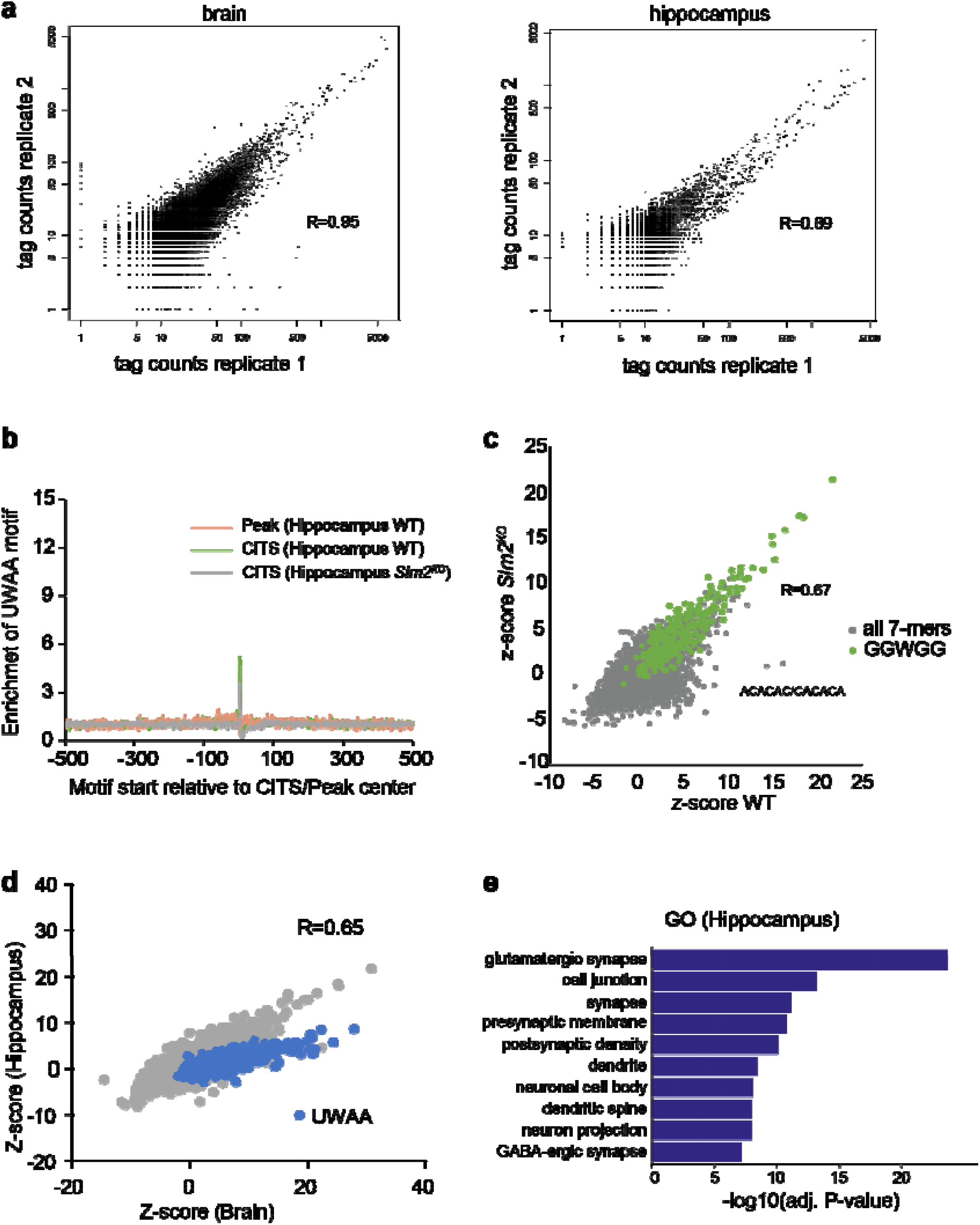
**a**, Correlation plot of tag numbers over called eCLIP tag clusters in replicate 1 (x-axis) and replicate 2 (y-axis) of whole brain (left panel) and hippocampus (right panel) eCLIP data. Pearson’s correlation coefficient is shown. **b**, Enrichment of UWAA around CITS is calculated from the frequency of UWAA starting at each position relative to the inferred crosslink sites, normalized by the frequency of the element in flanking sequences in hippocampus eCLIP data from wild-type and *Slm2*^*KO*^ hippocampus. Enrichment of UWAA around the CLIP tag cluster peak center is shown for comparison. **c**, Correlation plot of 7mer enrichment z-scores from WT and *Slm2*^*KO*^ hippocampal eCLIP data. The GGWGG motif identified in hippocampal WT eCLIP samples (highlighted in green) is found to the similar extent in global SLM2 knock-out control samples. Pearson’s correlation coefficient is shown. **d**, Correlation of 7-mer enrichment z-scores of 100nt region around peak center from whole brain (x-axis) and hippocampus (y-axis) eCLIP data. 7-mers including UWAA are highlighted in blue. Pearson’s correlation coefficient is shown. **e**, Gene Ontology analysis (DAVID tools) of genes with SLM2 binding sites in hippocampal eCLIP data identified by CLIPper/IDR. Top 10 enriched gene ontology categories for cellular compartment are displayed.

**Figure S2.**
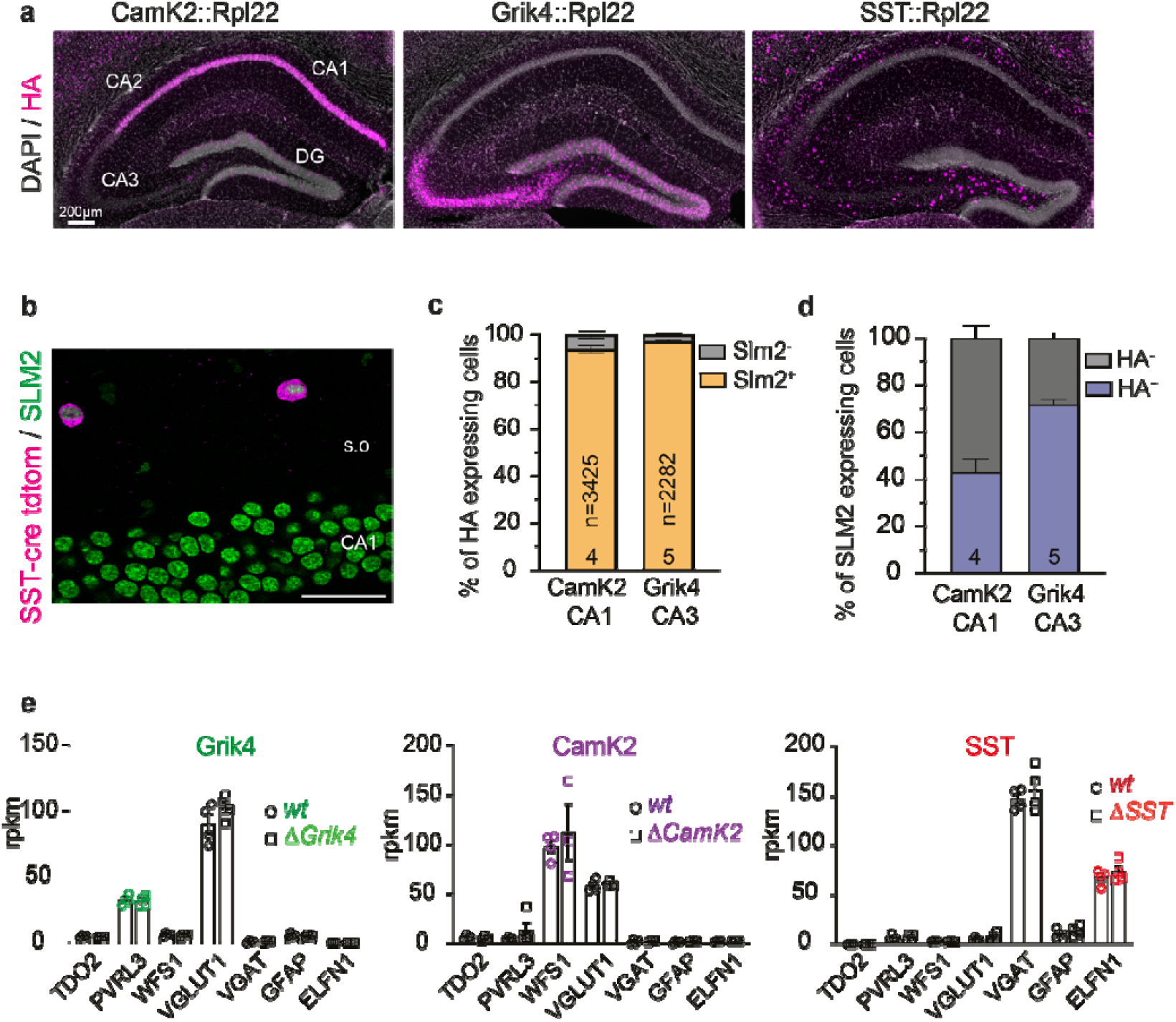
**a**, Representative images of cre-dependent expression of HA-tagged ribosomal protein L2 (Rpl22) in CA1 (CamK2::Rpl22), CA3 (Grik4::Rpl22) or SST+ interneurons (SST::Rpl22). Scale bar 200µm, DAPI (grey), HA (magenta). **b**, SLM2 (green) expression in CA1 pyramidal neurons and genetically marked SST+ interneurons (magenta) in the stratum oriens (s.o). Scale bar 40µm. **c**, Quantification of percentages of HA+ neurons defined by either CamK2 or Grik4 cre-recombinase which express SLM2 (SLM2+, orange). CamK2-cre: N=4 animals, n=3425 cells. Grik4-cre: N=5 animals, n=2282 cells. Mean of each replicate ± SEM. **d**, Quantification of SLM2+ neurons in either CA1 or CA3 layers which express Rpl22-HA (HA+, blue). Same images and numbers as for (c). Mean of each replicate ± SEM. **e**, Reads per kilobase million (rpkm) of cell class-specific marker genes in all analyzed cell types and individual replicates of *wt* and Δ*SLM2* animals (N=4). *TDO2*: DG marker, *PVRL3*: CA3, *RGS12*: CA2, *WFS1*: CA1, *VGLUT1*: excitatory neurons, *VGAT*: inhibitory neurons, *GFAP*: glia, *ELFN1*: SST neurons.

**Figure S3.**
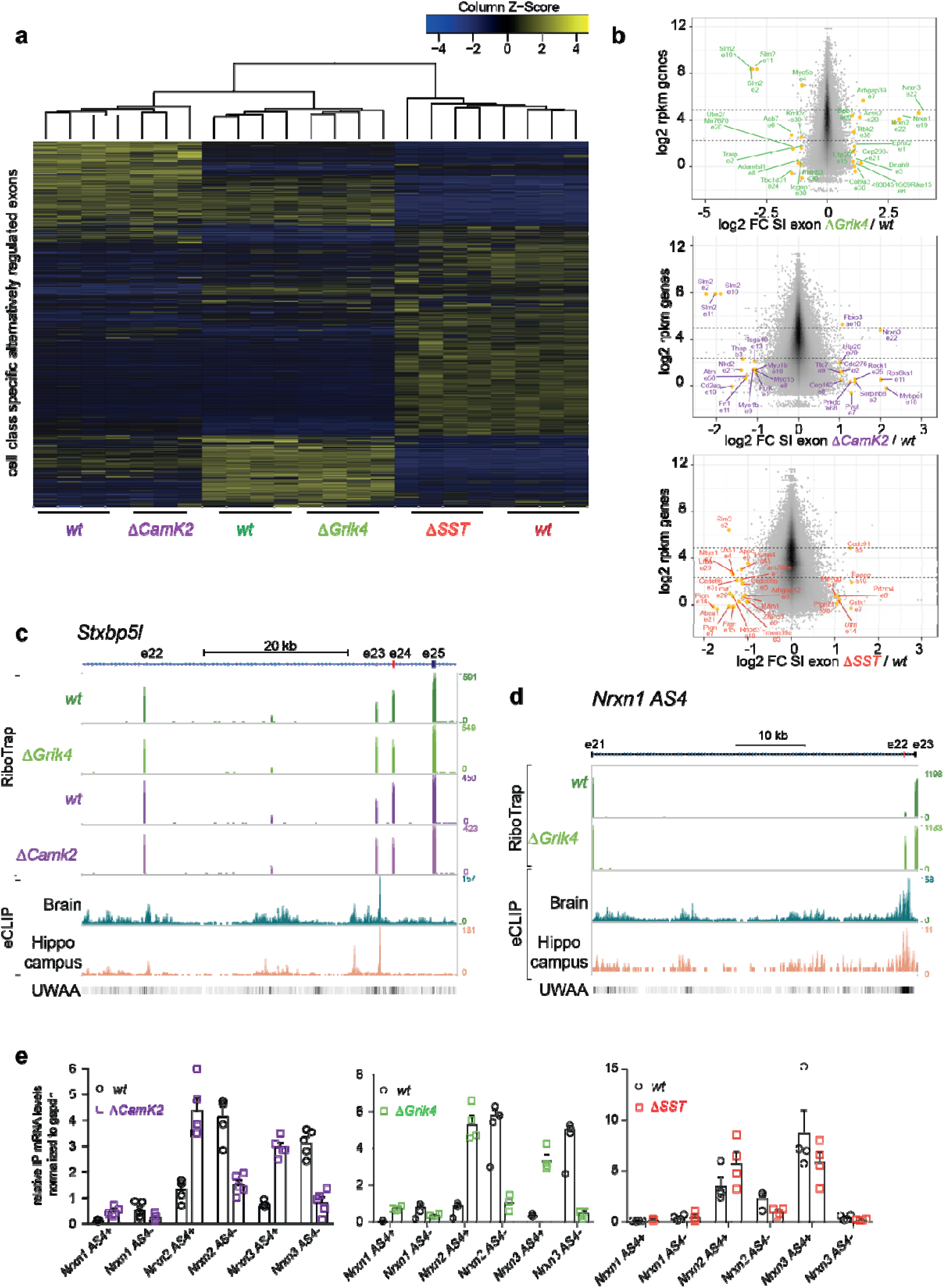
**a**, Heatmap of splicing indices (SI) of exons across individual replicates of *wt* and *Slm2* conditional mutants. Analyzed exons were defined by previously identified, cell class-specific alternative exons ^21^. Splicing indices were normalized by row and column. Detailed z-score values, gene and exon names are provided in Table S4. **b**, Log2 fold change SI (splicing index) and p-values for all detected exons (grey). Exons which are significantly differentially regulated by SLM2 in each cell class (called by exon analysis) are marked in yellow. **b**, Log2 fold change of SI of all detected exons (grey) and log2 rpkm values of the corresponding gene. Differentially regulated exons called by the exon analysis are marked in yellow. Gene names and exons involved in the splicing regulation are indicated in purple for differential changes in CamK2, green for Grik4 and red for SST. **c**,**d**, Integration of RiboTrap and eCLIP analysis for significantly de-regulated exons of *Stxbp5l* (c) and *Nrxn1* (d). SLM2 binding sites in the downstream introns and enrichment of the UWAA binding motif of the Grik4 comparison are illustrated. **e**, Quantitative PCR for alterations in *Nrxn* splicing at the alternatively spliced segment 4 (AS4). Relative *Gapdh* normalized mRNA levels of RiboTRAP IP samples in WT and ΔSLM2 samples. For all PCRs: *wt* CamK2 and Δ*CamK2*: N=5; *wt* Grik4 N=4 and Δ*Grik4* N=5, *wt* SST N=4 and Δ*SST* N=4, except for *Nrxn1*^*AS4+*^ N=3. *wt* CamK2 vs Δ*CamK2: Nrxn1*^*AS4-*^ p= 0.861, *Nrxn1*^*AS4+*^ p=0.0019, *Nrxn2*^*AS4-*^ p*=*0.0002, *Nrxn 2*^*AS4+*^ p=0.0003, *Nrxn3*^*AS4-*^ p*<0*.*0001, Nrxn 3*^*AS4+*^ p<0.0001; *wt* Grik4 vs Δ*Grik4 Nrxn1*^*AS4-*^ p=0.0035, *Nrxn1*^*AS4+*^ p<0.0001, *Nrxn2*^*AS4-*^ p<0.0001, *Nrxn2*^*AS4+*^ p<0.0001, *Nrxn3*^*AS4-*^ p<0.0001, *Nrxn3*^*AS4+*^ p<0.0001, *wt* SST vs Δ*SST Nrxn2*^*AS4-*^ p=0.0251.

**Figure S4.**
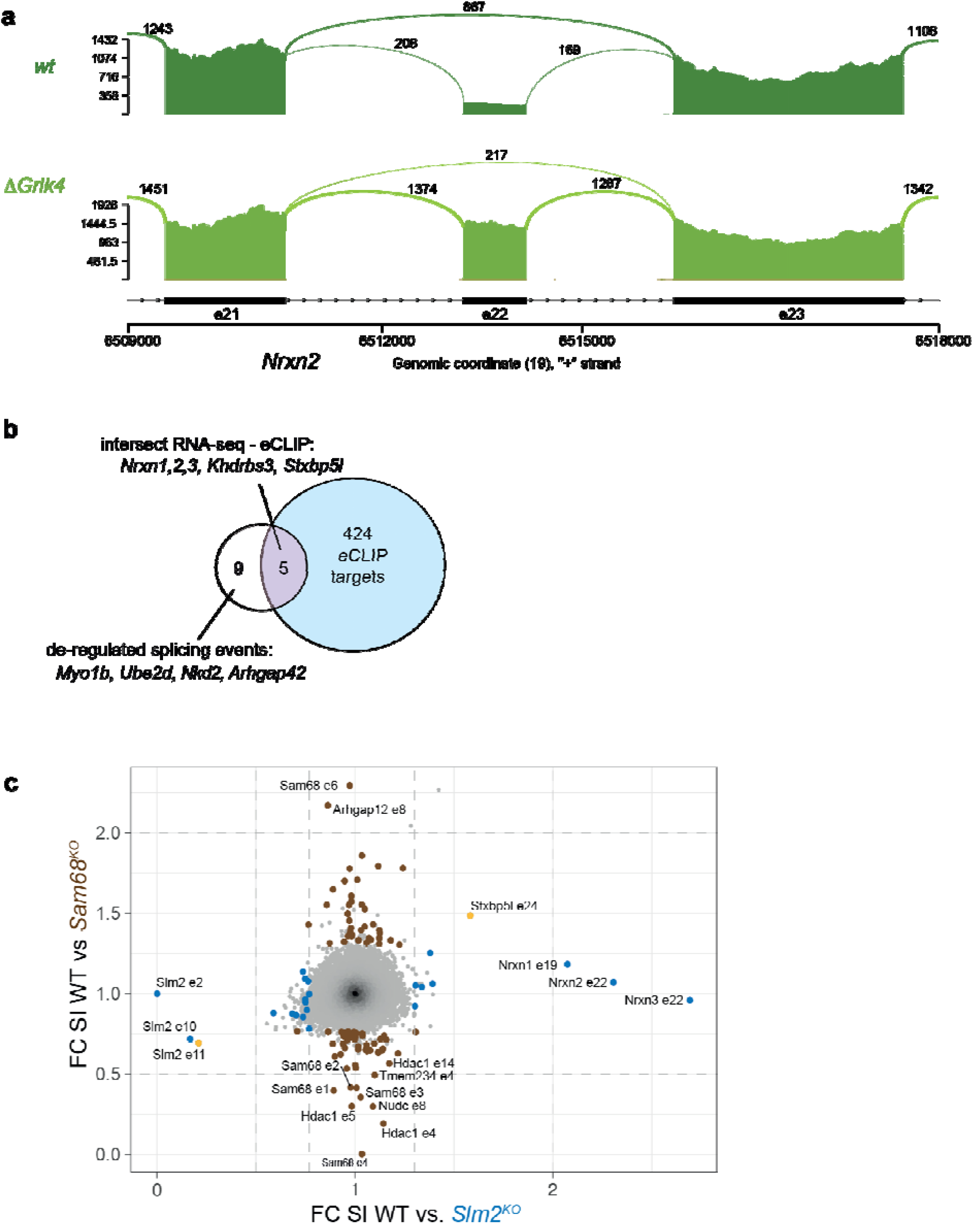
**a**, Representative sashimi plots illustrating read distribution and splice junctions on the *Nrxn2* gene at *AS4* for *wt* and Δ*Grik4* conditional mutant. Genomic coordinates and exon numbers are indicated below. Junction reads for exon-exon boundaries are noted and illustrated by line thickness. **b**, Venn diagram demonstrating the number of genes identified by eCLIP/IDR as bound (424), differentially alternatively spliced in *Slm2*^*KO*^ (9) or both bound and alternatively spliced (5). **c**, Correlation plot of the splicing index fold change (FC SI) in mouse hippocampus between WT and *Slm2* global knock-out (*Slm2*^*KO*^; ^37^) and WT and Sam68 global knock-out (*Sam68*^*KO*^; ^69^) for all detected exons (grey). Significantly differentially regulated exons (FC ± 30%, p-value 0.01) are marked in brown for *Sam68* and blue for *Slm2* mutants. Two exons, marked in yellow, are commonly de-regulated suggesting very little overlap in splicing regulation by SAM68 and SLM2 proteins.

**Figure S5.**
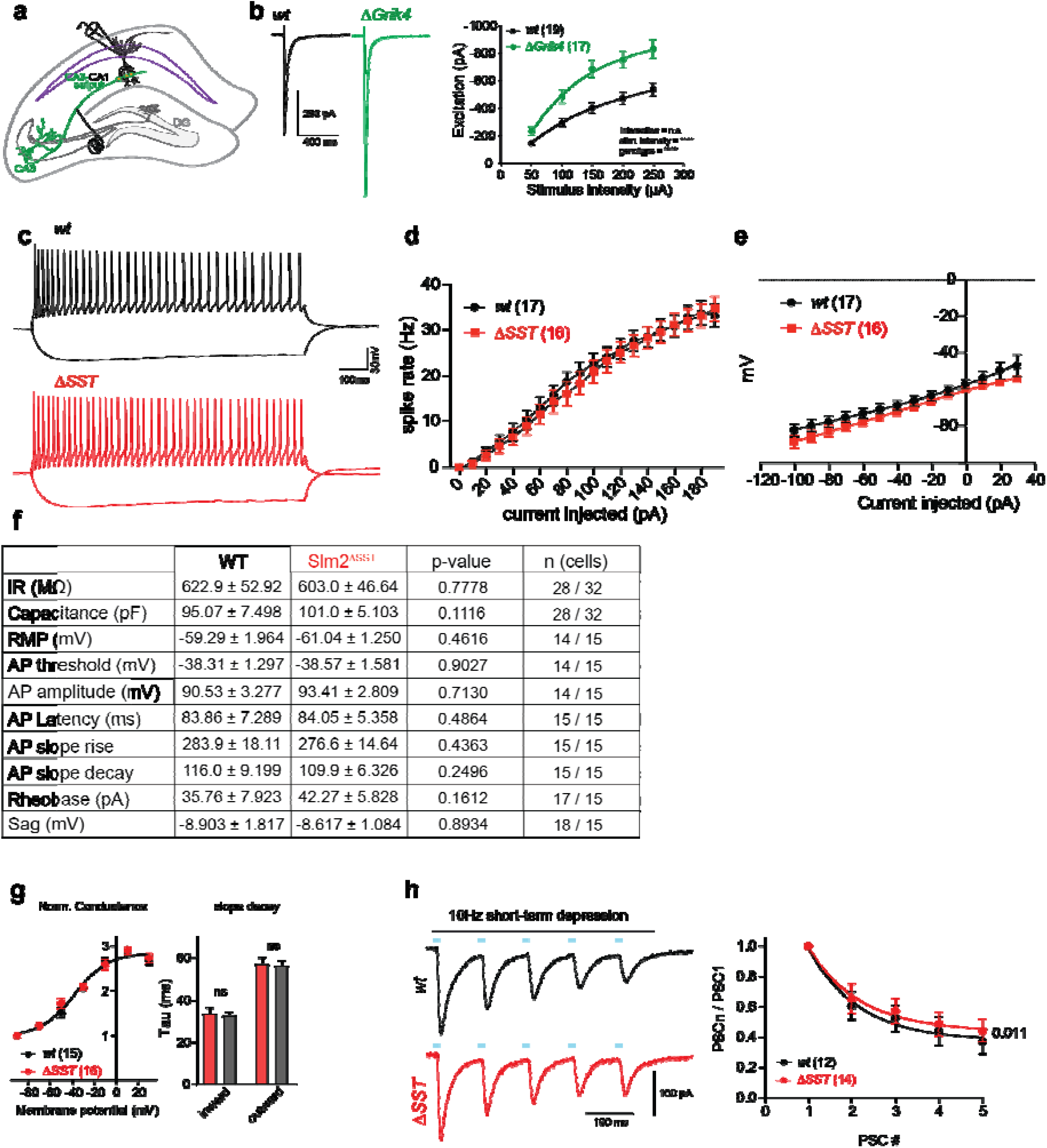
**a**, Experimental design for electrical stimulation of Schaffer collaterals and voltage clamp recordings in CA1 pyramidal cells in *wt* and Δ*Grik4* mutants. **b**, Representative traces of post-synaptic EPSCs *wt* (black) and Δ*Grik4* mutants (green). Electrically evoked EPSCs with various stimulation intensities in *wt* (n=19) and Δ*Grik4* mutants (n=17). Mean SD is displayed, two-way ANOVA was used for statistical analysis. **c**, Representative current clamp recordings to measure spike frequency of *wt* (black) and Δ*SST* (red) SST+ interneurons in s.o. Responses to a single 1s long -100pA or +150pA current injection. **d**, Frequency of action potential firing in response to increasing current injections. *wt* n= 17, Δ*SST* n=16 **e**, Analysis of changes in membrane potentials with increasing current in pA. The resting membrane potential is displayed at 0pA injection. *wt* n= 17, Δ*SST* n=16 **f**, Summary table of intrinsic electrophysiological properties of *wt* and Δ*SST* neurons. Mean ± SEM, p-values were determined by the corresponding t-tests based on assessment of normality distribution and standard deviation (see methods for details). **g**, left, a plot of normalized conductance versus membrane potential shows a clear voltage-dependence *wt* and Δ*SST* mutants. There was no genotype-dependent difference (p=0.60, Extra sum-of-squares F test). Right, analysis of decay times, weighted tau in milliseconds, of inward (at -90mV) and outward (at -10mV) currents showed no difference between genotypes (*wt* n=16 and Δ*SST* n=15) **h**, Example traces of IPSCs during repetitive stimulation at 10Hz, *wt* (black) and Δ*SST* (red). Group data of IPSCs normalized to the first peak. Mean ± SEM, *wt* n=14 and Δ*SST* n=12, Extra sum-of-squares F test for comparison of independent fits.

**Figure S6.**
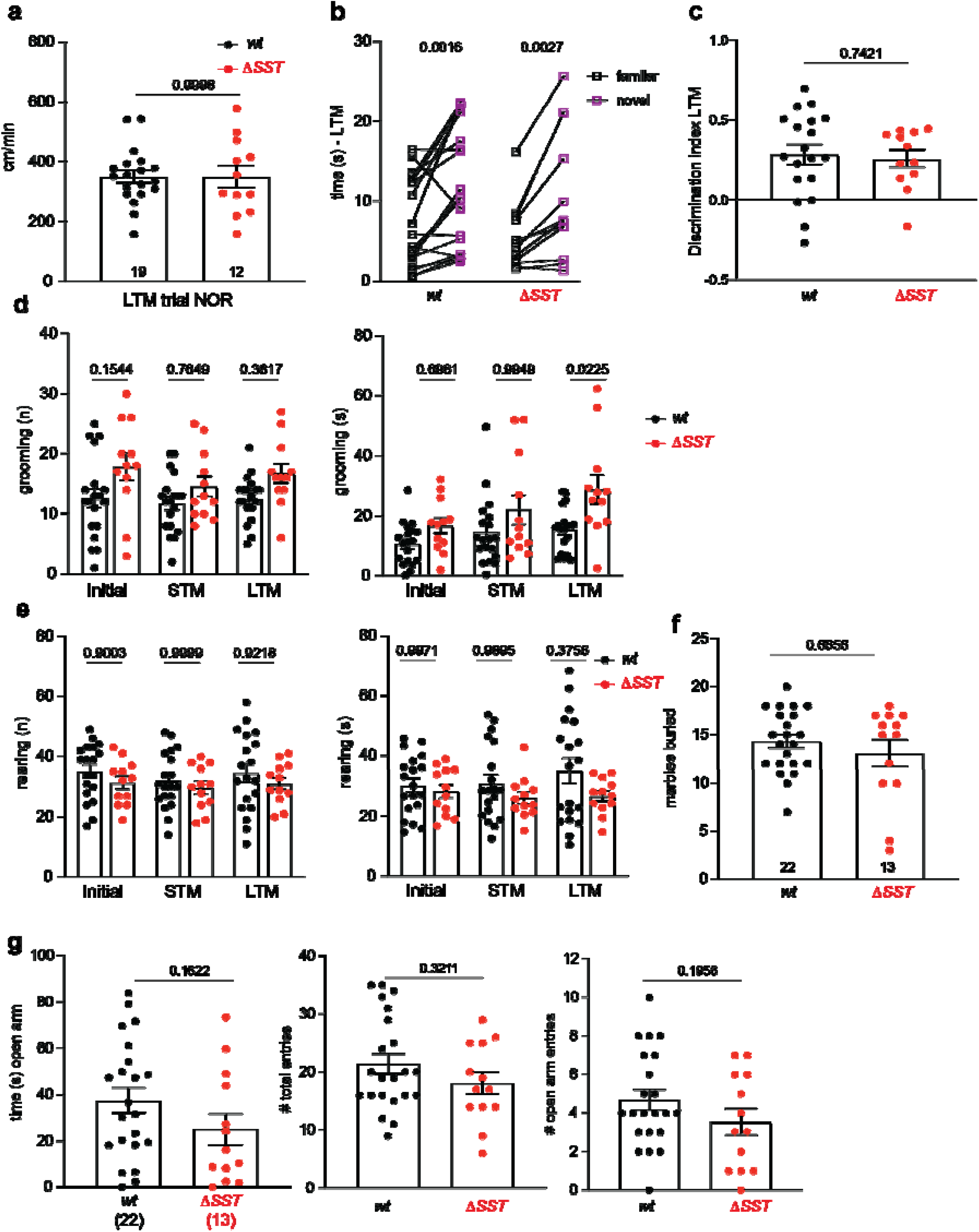
**a**, Quantification of velocity (cm/min) of mice during the long-term memory (LTM) test phases of the novel object recognition (NOR) task. Animal numbers for each task are indicated, Mean ± SEM, Unpaired t-test. **b**,**c**, Interaction time (in seconds) that mice spend with either a familiar (black) or novel (purple) object during a 5-min long-term memory trial (paired t-test) and discrimination index (unpaired t-test with Welch’s correction) are displayed. Mean ± SEM, *wt* n=19 and Δ*SST* n=12. **d**, Quantification of the number (left) and duration (right) grooming events of mice during the Open Field and phases of the NOR task. Mean ± SEM, One-Way ANOVA with Tukey’s multiple comparisons test. **e**, Quantification of the number (left) and duration (right) rearing events of mice during the Open Field and phases of the NOR task. Mean ± SEM, One-Way ANOVA with Tukey’s multiple comparisons test. **f**, Number of marbles buried when mice are placed in a novel homecage including 20 black marbles for 30min. Mean ± SEM, Mann Whitney t-test. *wt* n=22 and Δ*SST* n=13. **g**, Analysis of the amount of time mice spend in an open arm of the elevated plus maze during a 5min trial (left), their number of entries into either the open or closed arm (middle) and number of entries into the open arm (right). Mean ± SEM, upaired t-test. *wt* n=22 and Δ*SST* n=13.

**Figure S7.**
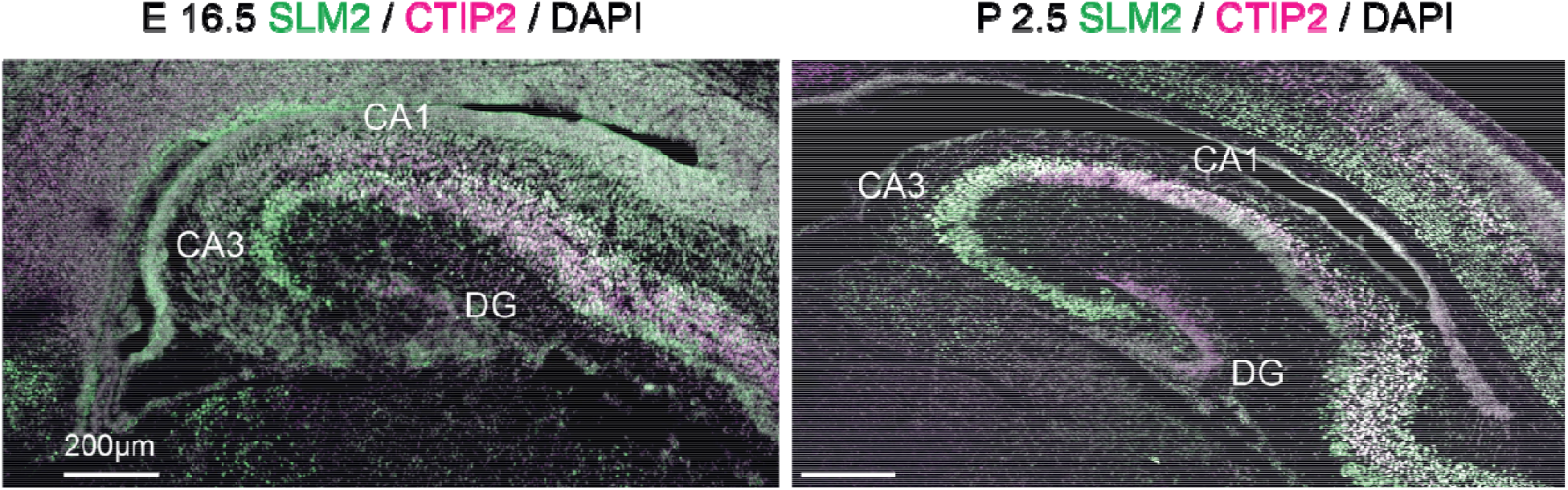
Hippocampal tissue from mice immunostained for SLM2 (green), CTIP2 (purple) and DAPI (grey) at E16.5 and P2.5. This demonstrates selective expression of SLM2 in CA1 and CA3 but not dentate granule cells at early stages of hippocampal development. Scale bar is 200µm.

### Supplementary Tables

**Table S1**. eCLIP data analysis for SLM2-bound mRNAs in mouse brain and hippocampal tissue. Tabs list annotation of all clusters identified in whole brain and hippocampus, respectively, called using the CTK pipeline ^44^.

**Tabl2 S2**. eCLIP data analysis for SLM2-bound mRNAs in mouse whole brain (WB) and hippocampal (HC) tissue analyzed with the CLIPper /IDR pipeline.

**Table S3**. Information on sequencing statistics. Global transcriptome analysis of hippocampal wild type and *Slm2* conditional knockout RiboTrap samples.

**Table S4**. Cell type specific alternative exons differentially regulated between wild type CA1, CA3 and SST interneurons in the hippocampus.

**Table S5**. Alternative exons differentially expressed between *Slm2* control and *Slm2* conditional knockout RiboTrap samples from CA1, CA3 and SST interneurons in the hippocampus.

**Table S6**. Differentially regulated patterns of alternative splicing in *Slm2* control and *Slm2* conditional knockout RiboTrap samples from CA1, CA3 and SST interneurons in the hippocampus.

## METHODS

### Mice

All procedures involving animals were approved by and performed in accordance with the guidelines of the Kantonales Veterinäramt Basel-Stadt. *Slm2* floxed mice have been generated in the Scheiffele laboratory and were previously described ^36^. *Rpl22-HA* (RiboTag) mice ^49^, *SST-cre* mice ^70^, Ai9 tdTomato ^71^, *CamK2-cre* mice ^72^, *Grik4-cre* mice ^73^, ChR2-flox mice ^74^ were obtained from Jackson Laboratories (Jax stock no: 011029, 013044, 007905, 005359, 006474, 012569 respectively). All lines were maintained on a C57Bl6/J background. The specificity of cre-lines for recombination of the Rpl22-allele and/or *Slm2*^*flox*^ was confirmed by immunohistochemistry. *Grik4-cre* mice require particular attention due to high rate of spontaneous germline recombination..

### Antibodies

Polyclonal antibodies for SLM2 and SAM68 were previously described ^35^. Additional antibodies are rat anti-HA (Roche, #11867431001, 1:1000), mouse anti-NeuN (Chemicon #MAB377 1:2000), and rabbit anti-CTIP2 (Novus Biologicals, #NB100-2600). Secondary antibodies included donkey anti-rat IgG-Cy3 and Cy5 (Jackson ImmunoResearch, 712-165-153, 706-175-148 1:1000). DAPI nuclear stain was co-applied with secondary antibody at a final concentration of 0.5 µg/ml.

### Primary hippocampal cell culture

Primary hippocampal culture was prepared from RjOrl:SWISS E16 mouse embryos. Hippocampi were dissected in plain DMEM (Invitrogen), minced and transferred in 2mL DMEM to a 15mL tube. 2mL of the 0.25% to the final concentration of 0.125% were added and incubated for 25min at 37°C in a water bath. Then 100µl of 1 mg/mL DNaseI (Roche) were added and incubated for additional 5min. The digestion solution was removed and hippocampi pieces were washed twice with DMEM containing 10% fetal bovine serum. Subsequently hippocampi pieces were triturated in 1mL neurobasal medium supplemented with B27 (Invitrogen), Glutamax (Invitrogen) and penicillin/streptomycin (Invitrogen). After trituration 4mL of neurobasal medium was added, cell suspension was filtered through a 70µl strainer and centrifuged for 10min at 100xg. Supernatant was removed and cell pellet was resuspended in 5ml neurobasal medium. 100.000 – 120.000 cells per well were plated in a 24 well plate with cover slips coated with poly-D-lysine and laminin.

### Immunohistochemistry, image acquisition and statistical analysis

Cultured cells were fixed at day-in-vitro (DIV) 12 with 4% PFA in 1x PBS for 10min at RT and washed 3x with 1xPBS. Cells were stained for endogenous Sam68 and SLM2 with polyclonal antibodies as previously described in Iijima et al., 2014. Briefly, fixed cells were blocked in blocking solution (5% milk, 0.1% Triton-X100 in PBS) for 1h at RT and then incubated with the primary antibodies anti-SLM2 (1:4000) and anti-SAM68 (1:2000) in blocking solution overnight at 4°C. After 3x washes with PBS cells were incubated for 1h with anti-rabbit-Cy3 and anti-guinea pig Cy5 antibodies in blocking buffer at RT, washed, stained with DAPI and mounted on glass slides. Images were acquired on an inverted LSM880 confocal microscope (Zeiss) using 63× Apochromat objectives in super-resolution Airyscan mode.

For immunohistochemistry on brain sections, postnatal animals (male and female) were transcardially perfused with fixative (4% paraformaldehyde in 100mM phosphate buffer, pH 7.2) and post-fixed over night at 4°. Embryonic brains were drop-fixed for 24h. Brain samples were immersed in 30% sucrose in 1X PBS for 48h, cryoprotected with Tissue-Tek optimum cutting temperature (OCT) and frozen at -80°. Early postnatal, adolescent and adult tissue was sectioned at 40µm on a cryostat and collected in 1X PBS, whereas embryonic tissue was sectioned at 20µm and collected directly on glass slides. Floating sections were blocked for 1h at RT in 10% Normal Donkey Serum + 0.05% Triton-X100, immunostained over night at 4°C with primary antibody incubation diluted in 10% Normal Donkey Serum + 0.05% Triton-X100. On slide staining was performed with blocking in 5% Normal Donkey Serum + 3% Bovine Serum Albumin + 0.05% Triton-X100 for 1h at RT, followed by RT incubation of primary antibody diluted in 1% Normal Donkey serum for 36h. Secondary antibodies were diluted in 1X PBS + 0.05% Triton-X100 for 2h at RT for protein detection, except for on slide staining for which antibodies were diluted in 1X Normal Donkey Serum for 1.5h at RT. Sections were mounted on glass slides with Prolong Diamond Antifade Mountant or Dako Fluorescence Mounting medium. Images were acquired at room temperature on an upright LSM700 confocal microscope (Zeiss) using 40x Apochromat objectives controlled by Zen 2010 software (1µm z-stacks). Hippocampal overview images were generated at room temperature on a Slidescanner AxioScan.Z1 (Zeiss) using a 10X objective. Stacks of 24µm thickness (4µm intervals) were used for a maximum intensity projection. Overview images of embryonic and P2 animals were taken on an LSM700 upright using a 10X Apochromat objective controlled by Zen 2010 software and image tiling. Images were analyzed and assembled using ImageJ (Fiji) and Adobe Illustrator software.

SLM2 intensity levels were characterized in NeuN+ cells residing in either CA1, CA3 or DG regions of the hippocampus. SST+ neurons were identified by genetic labelling using SST-cre mice crossed with tdTomato. Intensity levels were determined using in three dimensions using Imaris 7.0.0, Bitplane AG). Three dimensional surfaces were created around each nucleus of either cell class and the labelling intensity was automatically generated by the software based on the intensities of isolated pixels (determined as arbitrary units).

SLM2 knock-out efficiency was determined by comparing WT and SLM2 conditional mutants in either CA1 (Camk2), CA3 (Grik4) or the stratum oriens above CA1 for SST. Intensity levels for calling a neuron SLM2+ or SLM2-were previously determined by the intensity levels of SLM2 observed in the dentate gyrus of the same section. Following this, the number of SLM2+ and SLM2-neurons in the respective area imaged with 40x was quantified. For quantification in CA1 and CA3 mice were 5-6 weeks of age whereas quantification in tdTomato+, SST+ neurons was performed at p28. This strategy had been used as CamK2- and Grik4-cre recombinases are expressed at later developmental stages.

Quantification of the percentage of either SLM2 or HA+ cells was performed as follows: Within an area of either CA1 or CA3 of mice expressing Rpl22, the number of HA+ cells co-expressing SLM2 were determined. From the same image the number of SLM2+ cells that did or did not co-label with HA were additionally determined.

### eCLIP library preparation

The CLIP experiments were performed according to the eCLIP protocol from Nostrand et al. ^75^ with some modifications. Mouse whole brains or hippocampi were rapidly dissected on ice and immediately flash frozen in liquid nitrogen. The brain samples were ground on dry ice first in a custom-made metal grinder and a porcelain mortar. The frozen powder was transferred into a plastic Petri dish (10 or 6cm diameter) and distributed in a thin layer. Samples were UV-crosslinked 3x with 400mJ/cm^2^ on dry ice with a UV-crosslinker (Cleaver Scientific) with mixing and redistributing of the powder between single UV exposures. The crosslinked powder was re-suspended in 10ml (for 1 x whole brain) or 4.5ml (for 4 hippocampi) of the CLIP-lysis buffer (50mM Tris-HCl pH 7.5, 100mM NaCl, 1% NP-40, 0.1% SDS, 0.5% sodium deoxycholate) supplemented with 1 tablet per 10ml buffer of the protease inhibitors (Roche) and 4U per ml buffer Turbo-DNase (Thermofisher), transferred into a glass homogenizer and homogenized by 30 strokes on ice. 1ml aliquots were transferred to 2ml tubes, 10µl of RNaseI (Thermofisher) diluted in PBS (1:5 or 1:25) were added to a 1ml aliquot. Samples were incubated at 37°C with shaking (1’200 x rpm) for 5 min and put on ice. 10µl RNasin RNase-inhibitor (40U/l, Promega) were added, samples were mixed and centrifuged for 15min at 16’000 x g, 4°C. After centrifugation the supernatants were transferred to a new tube and 60µl from each sample were taken for sized matched INPUT (SMIn). To the rest 1ug/ml of affinity purified anti-SLM2 antibody was added and samples were incubated for 2h at 4°C in an overhead shaker. Then 10µl of Protein-A magnetic beads (Thermofisher) per 1µg antibody were added to each sample and samples were incubated for additional 2h at 4°C in an overhead shaker. Beads were washed 2x with the high salt wash buffer (50mM Tris-HCl pH7.5, 1M NaCl, 1mM EDTA, 1% NP-40, 0.1% SDS, 0.5% sodium deoxycholate), 2x CLIP-lysis buffer, 2x with low salt wash buffer (20mM Tris-HCl pH7.5, 10mM MgCl_2_, 0.2% Tween-20) and 1x with PNK buffer 70mM Tris-HCl pH6.5, 10mM MgCl_2_). Beads were re-suspended in 100µl PNK-mix (70mM Tris-HCl pH6.5, 10mM MgCl_2_, 1mM DTT, 100U RNasin, 1U TurboDNase, 25U Polynucleotide-Kinase (NEB)) and incubated for 20min at 37°C in a thermomixer with shaking (1200 x rpm). After RNA dephosphorylation beads were washed as before with 2x high salt, 2x lysis and 2x low salt buffers and additionally with 1x Ligase buffer (50mM Tris-HCl pH7.5, 10mM MgCl_2_). Beads were re-suspended in 50µl ligase mix (50mM Tris-HCl pH7.5, 10mM MgCl_2_, 1mM ATP, 3 % DMSO, 15% PEG8000, 30U RNasin, 75U T4 RNA-ligase (NEB)). 10µl of the beads / ligase mix were transferred to a new tube and 1µl of pCp-Biotin (Jena Bioscience) were added to validate IP of the RNA-protein-complexes by western blot. To the rest (40µl) 4µl of the RNA-adaptor mix containing 40µM of each RNA_X1A & RNA_X1B (IDT) were added and samples were incubated for 2h at RT. After adaptor ligation samples were washed 2x with high salt, 2x with lysis and 1x with low salt buffers. Beads were re-suspended in 1x LDS sample buffer (Thermofisher) supplemented with 10 uM DTT and incubated at 65°C for 10min with shaking at 1200 x rpm. Eluates or inputs were loaded on 4-12% Bis-Tris, 10-well, 1.5mm gel (Thermofisher) and separated at 130V for ca. 1.5h. Proteins were transferred to the nitrocellulose membrane (Amersham) overnight at 30V. After transfer the membranes were placed in a 15cm Petri dish on ice and an area between 55 and 145kDa was cut out in small pieces and transferred to 2ml tube. For CLIP samples RNA extraction, reverse transcription using AR17 primer, cDNA clean-up using silane beads (Thermofisher), second adaptor ligation (rand103Tr3) and final cDNA purification were performed as previously described ^75^. For sized matched input samples (SMIn) isolated RNA was dephosphorylated.

The sequencing libraries were amplified using Q5-DNA polymerase (NEB) and i50X/i70X Illumina indexing primers (IDT). 14 cycles were used for the amplification of whole brain libraries and 16 cycles for hippocampus libraries. Corresponding SMIn libraries were amplified with 12 cycles for whole brain and 16 cycle for hippocampus samples. The amplified libraries were purified and concentrated first with ProNEX size selective purification system (Promega) using sample/beads ratio of 1/2.4. The purified libraries were loaded on a 2% agarose gel, the area corresponding to the size between 175bp and 350bp was cut and the libraries were extracted from the gel using gel extraction kit (Machery&Nagel) and eluted with 16µl.

Concentrations and size distributions of the libraries were determined on Fragment analyzer system (Agilent). 75bp paired-end sequencing was performed on the NextSeq500 platform using Mid Output Kit v2.5 (150 cycles). Adaptor and primer sequences used in this study are listed in the Key Resources Table.

### eCLIP data processing

The raw reads were processed to obtain unique CLIP tags mapped to mm10 using CTK ^42^, as described previously ^44^. Unique tags from replicates were combined for all analyses. Significant CLIP tag clusters were called by requiring P<0.05 after Bonferroni multiple-test correction. Crosslinking-induced truncation sites (CITS) were called by requiring FDR<0.001. We performed 7-mer enrichment analysis using significant peaks with peak height (PH)≥10 tags. Peaks were extended for 50 nt on both sides relative to the center of the peak to extract the foreground sequences. Background sequences were extracted from the flanking regions of the same size (−550, -450) and (450, 550) relative to the peak center. Sequences with more than 20% of nucleotides overlapping with repeat masked regions were discarded. 7-mers were counted in repeat-masked foreground and background sequences, and the enrichment of each 7-mer in the foreground relative to the background was evaluated using a binomial test, from which z-score (and a p-value) was derived. In parallel, eCLIP data was analyzed following the ENCODE pipeline using the CLIPper peak calling tool (https://github.com/YeoLab/clipper; https://github.com/YeoLab/eclip), followed by IDR (https://github.com/nboley/idr) to identify peaks reproducibly identified between replicates. Both analysis pipelines (CTK and CLIPper) gave similar results and led to the same biological conclusions.

Data was deposited at GEO: GSE220062

### Motif analysis

UWAA motif sits were searched in genic regions and their conservation was evaluated using branch length score (BLS) estimated from multiple alignments of 40 mammalian species ^76^. *De novo* motif discovery was performed using mCross, an algorithm that augments the standard PWM model by jointly modeling RBP sequence specificity and the precise protein-RNA crosslink sites at specific motif positions at single-nucleotide resolution ^44^. For this analysis, the top 10 enriched 7-mers from significant peak regions were used as seed to search for overrepresented motifs around CITS sites, as described previously ^44^. The motif with the maximum motif scores was chosen as the represented motif.

### Gene Ontology analysis

Gene Ontology was performed using the DAVID functional annotation (cellular compartment) online tool (https://david.ncifcrf.gov/tools.jsp). The input list for the background was the list of all genes detected in a hippocampal sample analyzed by bulk RNA sequencing ^37^. Genes that had significant peak expression in the CLIP dataset either for hippocampus or whole brain samples were used. The top ten significant (Benjamini Hochberg for p-value correction), and non-redundant, terms were displayed in Figure 1g (whole brain) and Figure S1e (hippocampus).

### RiboTRAP pulldowns, RNA purification and quality control

RiboTRAP purifications were performed as previously described ^21,77^. For CamK2 and Grik4 pull downs animals were between postnatal day 39-42, for SST neurons between postnatal day 28-30. Mice were anesthesized with isoflurane and following cervical dislocation hippocampal tissue was rapidly dissected in ice-cold PBS and lysed in 0.5ml for single animals (Camk2 and Grik4) and 1mL for pools of two animals (SST) in homogenization buffer containing 100mM KCl, 50mM Tris-HCl pH 7.4, 12mM MgCl_2_, 100µg/mL cycloheximide (Sigma-Aldrich # 66-81-9), 1mg/mL heparin (Sigma-Aldrich #H339350KU), 1x complete mini, EDTA-*free* protease inhibitor cocktail (Roche #11836170001), 200 units/mL RNasin plus inhibitor (Promega #N2618) and 1mM DTT (Sigma-Aldrich #3483-12-3). The lysate was centrifuged for 10min at 2.000xg, 1% final concentration of Igepal-CA630 (Sigma Aldrich #18896) was added to the supernatant and incubated on ice for 5min, followed by an additional spin at 12.000xg. 1% of input was saved in RLTplus buffer (Qiagen RNeasy Micro Kit #74034) supplemented with 2-Mercaptoethanol before 20µl or 15µl of HA-magnetic beads (Pierce, #88837) were added to the excitatory or inhibitory pull down, respectively. Lysate/bead mixtures were incubated at 4° for 3-4hours under gentle rotation and were afterwards washed 4 times with wash buffer containing 300mM KCl, 1% Igepal-CA630, 50mM Tris-HCl, pH7,4, 12mM MgCl_2_, 100 µg/mL Cycloheximide and 1mM DTT. RNA was eluted from beads with 350µl RLT plus buffer supplemented with 2-Mercaptoethanol as per manufacturers instructions.

RNA of input and RiboTrap IP samples was purified using the RNeasy Plus Micro Kit (Qiagen #74034) following manufacturer’s instructions. RNA was further analyzed using an RNA 6000 Pico Chip (Agilent, 5067-1513) on a Bioanalyzer instrument (Agilent Technologies) and only RNA with an integrity number higher than 7.5 was used for further analysis. RNA concentration was determined by Fluorometry using the QuantiFluor RNA System (Promega #E3310) and 20ng of RNA was reverse transcribed for analysis of marker enrichment by quantitative PCR. Only samples which had an enrichment for hippocampal layer specific excitatory neuron markers and a de-enrichment for inhibitory or glia markers were further used for CamK2 and Grik4. SST pulldowns exhibited an enrichment in inhibitory neuron markers and a de-enrichment in excitatory and glia markers.

### Library preparation and illumina sequencing

Four biological replicates per cell class and genotype were further analyzed. Library preparation was performed with 50ng of RNA using the TruSeq PolyA+ Stranded mRNA Library Prep Kit High Throughput (Illumina, RS-122-2103). Libraries were quality-checked on a Fragment Analyzer (Advanced Analytical) using the Standard Sensitivity NGS Fragment Analysis Kit (Advanced Analytica, DNF-473), revealing high quality of libraries (average concentration was 49±14 nmol/L and average library size was 329±8 base pairs). All samples were pooled to equal molarity and the pool was quantified by PicoGreen Fluorometric measurement. The pool was adjusted to 10pM for clustering on C-Bot (Illumina) and then sequenced Paired-End 101 bases using the HiSeq SBS Kit v4 (Illumina, FC-401-4003) on a HiSeq 2500 system. Primary data analysis was performed with the Illumina RTA version 1.18.66.3 and bcl2fastq-v2.20.0.422.

### Quality control and RNA-seq pre-processing

The gene expression and alternative splicing analysis of the RNA-Sequencing data were performed by GenoSplice technology (www.genosplice.com) and have been additionally described in ^21^. Data quality, reads repartition (e.g., for potential ribosomal contamination), and insert size estimation were performed using FastQC v0.11.8, Picard-Tools v1.119, Samtools 1.13 and rseqc v2.3.9. This first quality check identified one sample in the pool of DCamK2 which displayed an accumulation of reads on the 3’end and displayed higher ribosomal contamination. Thus, this sample was excluded from further analyses. Reads were mapped using STARv2.4.0 ^78^ on the mm10 Mouse genome assembly. Reads were mapped using STARv2.4.0 ^78^ against the exons defined in the proprietary Mouse FAST DB v2016_1 database ^79^, using a mismatch cutoff of 2 and discarding reads with 10 or more alignments. The minimum chimeric segment length was 15. Read counts were summarized using featureCounts ^80^ in two stages. First, unique reads per exon were counted. In the second stage, multimapping reads were fractionally allocated to exons based on the distribution of unique counts of exons within a gene. Total counts were then calculated based on three constitutivity classes defined in FAST DB: class 2 includes exons present in more than 75% of annotated transcripts for a gene (“constitutive”), class 1 includes exons present in 50-75% of transcripts (“semi-constitutive”), and class 0 includes exons present in less than 50% of transcripts (“alternative”). Total counts per gene were summed from constitutivity class 2 exons if their FPKM values exceed 96% of the background FPKM based on intergenic regions. If counts from class 2 exons were insufficient to exceed the detection threshold, class 1 and eventually class 0 exon counts were included to reach the detection threshold.

Data was deposited at GEO: GSE209870

### Differential gene expression analysis

Differential regulation of gene expression was performed as described ^81^. Briefly, for each gene present in the proprietary Mouse FAST DB v2016_1 annotations, reads aligning on constitutive exons of the gene are counted. Based on these read counts, normalization and differential gene expression are performed using DESEq2 ^82^. Background expression was defined by reads aligning to intergenic regions, thus, only genes are considered as expressed if their RPKM value (reads per kilo base of transcript per million mapped reads) is greater than 96% of the background RPKM value based on intergenic regions. Only genes expressed in at least 3 out of 4 biological replicates for Grik4 and SST; and in at least 2 out of 3 biological replicates for CamK2 were further analyzed. For all expressed genes, DESeq2 values were generated (values were normalized by the total number of mapped reads of all samples). Fold change in gene expression was calculated by pairwise comparisons, comparing the normalized expression value in the respective WT condition to the corresponding ΔSLM2 condition and p-value (unpaired Student’s t-test) and adjusted p-value (Benjamin and Hochberg) were calculated. Results were considered significantly different for adjusted p-values ≤ 0.05 and fold changes ≥ ± 1.5.

### Alternative splicing analysis

Identification of alternatively spliced exons was performed with two analysis approaches as previously described ^21^: “**exon”** and “**pattern”** analysis. The exon analysis takes reads mapping to exonic regions and to exon-exon junctions into account. When reads map onto exon-exon junctions, the reads were assigned to both exons and the minimum number of nucleotides is 7 in order that a read is considered mapped to an exon. An exon was considered to be expressed if the FPKM value (Fragments per kilobase of transcript per million mapped reads) was greater than 96% of the background FPKM value based on intergenic regions. Only exons that were expressed in at least 3 out of 4 biological replicates for Grik4 and SST; and in at least 2 out of 3 biological replicates for CamK2 were further analyzed. Furthermore, for every expressed exon a splicing index (SI) was calculated: This is the ratio between read density on the exon of interest (=number of reads on the exon / exon length in nucleotides) and read density on constitutive exons of the same gene (with constitutive exons defined in FAST DB). The second type of alternative splicing analysis is the Pattern analysis. This type of analysis is taking known splicing patterns annotated in the FAST DB database into account ^79^. For each gene all annotated splicing patterns are defined and a SI is generated by comparing the normalized read density to the alternative annotated patterns.

The Log2 fold change (FC) and p-value (unpaired Student’s t-test) for both the exon and pattern analysis was calculated by pairwise comparisons of the respective SI values. Results were considered significantly different for p-values ≤ 0.01 and log2(FC) ≥1 or ≤-1. Sashimi plots were generated with Sashimi.py ^83^.

### qPCR analysis for alternative exon usage of *Nrxns* at AS4

Ribotag purified material was reverse transcribed and quantitative PCR was performed. qPCRs were performed on a StepOnePlus qPCR system (Applied Biosystems). Assays were used with a TagMan Master Mix (Applied Biosystems) and comparative C_T_ method. mRNA levels were normalized to the amount of *Gapdh* cDNA present in the same sample.

Custom gene expression assays were from Applied Biosystems and are described in ^35^.

### Electrophysiology

#### Slice preparation

Adult mice (P56-70) were anaesthetized with isoflurane (4% in O_2_, Vapor, Draeger) or with intraperitoneal injection of ketamine/xylazine (100mg/kg and 10mg/kg), and killed by decapitation, in accordance with national and institutional guidelines. For recordings in SST interneurons P17-18 animals were used. Slices were cut as previously described ^84^. Briefly, the brain was dissected in ice-cold sucrose-based solution at about 4 °C. Horizontal 300- to 350- μm-thick hippocampal brain slices were cut at an angle of about 20° to the dorsal surface of the brain along the dorso−ventral axes of the hippocampus using a Leica VT1200 vibratome. For cutting and storage, a sucrose-based solution was used, containing 87 mM NaCl, 25 mM NaHCO_3_, 2.5 mM KCl, 1.25 mM NaH_2_PO_4_, 75 mM sucrose, 0.5 mM CaCl_2_, 7 mM MgCl_2_ and 10 mM glucose or 10 mM dextrose (equilibrated with 95% O_2_/ 5% CO_2_). Some slices were prepared with additional 1-5 mM ascorbic acid and/or 3 mM pyruvic acid. Slices were kept at 32-35°C for 30 min after slicing and subsequently stored at room temperature either in cutting solution or in artificial cerebrospinal fluid (ACSF): 124mM NaCl, 2.5 mM KCl, 1.25 mM NaH_2_PO_4_, 2 mM CaCl_2_, 1-2 mM MgSO_4_, 26 mM NaHCO_3_, 10 mM dextrose or 10 mM glucose until experiments were performed at 21 to 22° C. For experiments, slices were, transferred to the recording chamber and perfused (1.5– 2.0 ml/min) with oxygenated ACSF at room temperature.

#### Whole-cell voltage-clamp recordings of EPSCs in CA1 pyramidal neurons

Hippocampal CA1 pyramidal neurons were visually identified in the pyramidal cell layer using Dodt-contrast video microscopy. Somatic whole-cell recordings were made from CA1 pyramidal neurons, which were voltage clamped with a Multiclamp 700B amplifier, and currents were digitized by Digidata 1440a. Patch pipettes (4–8 MΩ) were filled with voltage-clamp solution for excitation response curves: 125mM Cs-gluconate, 2 mM CsCl, 5 mM TEA-Cl, 4 mM ATP, 0.3 mM GTP, 10 mM phosphocreatine, 10 mM HEPES, 0.5 mM EGTA, and 3.5 mM QX-314. Data were filtered at 2 kHz, digitized at 10 kHz, and analyzed with Clampfit 10. SC afferents were stimulated with a small glass bipolar electrode prepared from theta glass (Sutter, BT-150-10) and passed once through a Kimwipe to make a 25-50µM opening. Excitation response curves were quantified from the average of the peak from ten evoked EPSCs (0.1Hz) voltage-clamped at -70mV – near the reversal potential for GABAAR-mediated inhibition. Short-term plasticity was induced with five stimuli of equal intensity at 40 Hz and voltage-clamped at -70mV. Data was analyzed with custom software written for this project using Python 3.7 and the pyABF module (http://swharden.com/pyabf). Significance was assessed by a two-way ANOVA for multiple comparisons.

#### Whole-cell voltage-clamp recordings of IPSCs in CA1 pyramidal neurons

CA1 pyramidal neurons were visually identified in the pyramidal cell layer using infrared differential interference contrast (IR-DIC) video microscopy. Patch-pipettes (2–4.5 MΩ) were filled with a Cs gluconate-based solution containing: 135mM CsGluc, 2mM CsCl, 10mM EGTA, 10mM Hepes, 2mM MgCl_2_, 2mM Na_2_ATP, 2mM TEA-Cl, 5mM QX314 adjusted to pH 7.3 with CsOH.

A diode laser (DL-473, Rapp Optoelectronic) was coupled to the epifluorescent port of the microscope (Zeiss Examiner, equipped with a 63× NA1.0 water immersion objective; Carl Zeiss Microscopy GmbH, Jena, Germany) via fiber optics. The laser was controlled via TTL pulses. For the optogenetic activation of the axon of SST+ interneurons, the field of view was shifted to stratum lacunosum moleculare and laser light was applied at intensities of 0.1-3.2 mW for 2 ms.

Optogenetically evoked IPSCs were recorded in presence of 25 µM AP5 and 10 µM NBQX. During the assessment of the voltage dependence of optogenetically activated GABA receptors, the series resistance was compensated at 80%. Membrane potentials were corrected offline by the calculated liquid junction potential of -15.7 mV ^85^.

#### Voltage- and current-clamp recordings in SST+ OLM interneurons

In slices from SST-Cre x Ai9tdTomato x SLM2^flox^ mice, putative OLM interneurons were visually identified according to their fluorescence, location in stratum oriens close to the alveus and by their morphology with an oval cell body and bipolar morphology oriented in parallel to the alveus. Somatic whole-cell recordings from s.o SST interneurons close to the alveus were clamped with a Multiclamp 700B amplifier (Molecular Devices, Sunnyvale, CA) and identified using epifluorescence microscopy. Signals were low-pass filtered at 2kHz, digitized at 10kHz. For voltage-clamp recordings, patch pipettes used were between 2-6 MΩ and filled with either with a solution containing 135mM CsMeSO_3_, 10mM Hepes, 9mM NaCl, 0.3mM EGTA, 4mM Mg-ATP, 0.3 Na-GTP, 5mM QX-314, 0.1mM Spermine, 303mOsm, pH=7.3 or 135mM CsGluc, 2mM CsCl, 10mM EGTA, 10mM Hepes, 2mM MgCl_2_, 2mM Na_2_ATP, 2mM TEA-Cl, 5mM QX314 adjusted to pH 7.3 with CsOH. Cells which had a change in series resistance ≥ 20% from start to the end of the experiment, or a series resistance higher than 25 were excluded from the analysis. Membrane resistance, series resistance and capacitance were constantly monitored by a -5mV step at the end of the trace.

To stimulate CA1 pyramidal neuron axon collaterals, the pipettes were placed into the border region between stratum oriens and alveus at a distance of approx. 200-250µM from the recorded neuron, and electrical stimulation was applied at low intensity (10-50 µA, at least 20x every 10s). The minimal first average response amplitude had to be at least 60pA in order to be further analyzed. 100 µM picrotoxin and 1 µM CGP54626 were added to block GABA_A_-mediated postsynaptic currents and GABA_B_ signaling, respectively.

Measurements of intrinsic properties were performed in current-clamp I_c_ with the following internal solution: 135mM K-gluconate, 5mM NaCl, 5mM MgATP, 0.3mM NaGTP, 10mM Phosphocreatine, 10mM Hepes without the addition of blockers.

*Data* was analyzed was performed offline using the open-source analysis software Stimfit ^86^ (https://neurodroid.github.io/stimfit) and customized scripts written in Python. The analysis of voltage-clamp data was performed on mean waveforms. Cumulative distribution analysis (in%) was performed in Prism. Amplitudes were analysed on individual events of every cell, whereas inter-event intervals were calculated based on the frequency of events per 10s sweep.

#### Drugs

All drugs were stored as aliquots at -20°C. D-AP5 (50 mM; Tocris) was dissolved in water. Picrotoxin was dissolved at 50 mM in ethanol. CGP 54626 hydrochloride (10 mM; Tocris) and NBQX (20 mM; Tocris) were dissolved in DMSO.

### Behavioral Analysis

Mice used for behavioral experiments were maintained in C57/Bl6J background, male, between 7-9 weeks of age and housed under standard laboratory conditions on a 12h light/dark cycle. All tests were carried out during the light cycle, with standard ceiling light and in at least 3 independent trials. All statistical data are mean ± SEM. Every animal was tested in all behavioral assays (battery testing).

#### Open Field

Mice were individually exposed to a square open field arena (50 × 50 × 30 cm) made of grey plastic for 10min. Velocity (cm/min) and time spent in the center were extracted from a video-based EthoVision10 system (Noldus).

#### Novel Object recognition task

Animals tested in the Open Field arena on the day before the experiment, were exposed to two identical objects (culture dish flask filled with sand) for 5min in the first trial (acquisition). After 1hour, we tested for Short-term memory by 5min exploration of one familiar (flask) and one novel object (Lego block). The time spent investigating the objects, sniffing less than a centimeter from or touching the object, was scored manually. The time mice spent on the objects was excluded (exploration not directed at the object itself). Only mice spending at least 2 seconds with the objects in total were included in the analysis. Calculation of discrimination ratio: (time spent with novel object – time spent with familiar object) / total time investigating both objects. Distance travelled was extracted from the video-based EthoVison10 system (Noldus). Time and number of grooming or rearing events, and the time spent investigating the objects was scored manually.

#### Elevated Plus Maze

Animals were placed in the center of the maze (arms are 35 cm x 6 cm and 74 cm above the ground) facing the closed arms. The time spent on the open arm was measured during the 5 min test. In addition, the number of total entries (open arms and closed arms) were counted manually.

#### Marble Burying

Animals were exposed to 20 identical black marbles distributed equally (4×5) in a standard Type II long cage with 5 cm high bedding for 30 min with ceiling light. For a marble to be counted as buried, approximately ≥ 75% of its area had to be below the bedding material.

## QUANTIFICATION AND STATISTICAL ANALYSIS

Quantification of electrophysiological data was performed using stimfit, histology and behavioral data was quantified by an experimenter blinded towards genotype. Statistical analysis for differential gene expression and alternative splicing events of RNA Sequencing experiments was performed in R and adjusted with the Benjamini Hochberg correction. All other statistical analysis was conducted using Prism version 8.0 and 9.0. Data was tested for normality with the Kolmogorov-Smirnov test and similar standard deviation before appropriate t-test were chosen for molecular, electrophysiological and behavioral experiments. Paired t-tests were applied for the comparison of interaction time between familiar and novel objects. When assessing changes in the electrophysiological or behavioral properties in which multiple groups were compared, one or two-way ANOVA’s with appropriate correction for multiple comparisons (either Šídák’s or Tukey’s multiple comparisons tests) were performed.

## Notes

### Competing Interest Statement

The authors have declared no competing interest.

